# Refining the Identity and Role of Kv4 Channels in Mouse Substantia Nigra Dopaminergic Neurons

**DOI:** 10.1101/2021.02.01.429100

**Authors:** Alexis Haddjeri-Hopkins, Mónica Tapia, Jorge Ramirez-Franco, Fabien Tell, Béatrice Marqueze-Pouey, Marianne Amalric, Jean-Marc Goaillard

**Affiliations:** UMR_S 1072, Aix Marseille University, INSERM, Faculté de Médecine Secteur Nord, 51 boulevard Pierre Dramard, 13015 Marseille, FRANCE; UMR7291, Aix Marseille University, CNRS, LNC, 3 place Victor Hugo, 13331 Marseille, FRANCE

**Author notes:** These authors contributed equally to this work (Alexis Haddjeri-Hopkins, Mónica Tapia). Corresponding author **Corresponding author**: Jean-Marc GOAILLARD. **Author contributions:** M.A., F.T. and J-M.G. designed research. A.H-H., M.T., J.R.F., F.T. and B.M-P. performed research. A.H-H., M.T., J.R.F., F.T. and J-M.G. analyzed data. A.H-H., M.T., J.R.F., F.T., M.A. and J-M.G. wrote the manuscript.

## Abstract

Substantia nigra pars compacta (SNc) dopaminergic (DA) neurons display a peculiar electrical phenotype characterized *in vitro* by a spontaneous tonic regular activity (pacemaking activity), a broad action potential and a biphasic post-inhibitory response. Several studies in rodents have underlined the central role played by the transient A-type current (I_A_) in the control of pacemaking activity and post-inhibitory rebound properties, thereby influencing both DA release and the physiological response of SNc neurons to incoming inhibitory inputs. Kv4.3 potassium channels were considered to be fully responsible for I_A_ in these neurons, their density being tightly related to pacemaking frequency. In spite of this crucial electrophysiological role, we show that Kv4.3^-/-^ transgenic mice exhibit minor alterations in locomotion and motor learning, although no compensation by functionally overlapping ion channels is observed in Kv4.3^-/-^ SNc DA neurons. Using antigen retrieval immunohistochemistry, we further demonstrate that Kv4.2 potassium channels are also expressed in SNc DA neurons, even though their contribution to I_A_ appears significant only in a minority of neurons (~5-10%). Using correlative analysis on recorded electrophysiological parameters and multi-compartment modeling, we then demonstrate that, rather than its conductance level, I_A_ gating kinetics (inactivation time constant) appear as the main biophysical property defining post-inhibitory rebound delay and pacemaking frequency. Moreover, we show that the hyperpolarization-activated current (I_H_) has an opposing and complementary influence on the same firing features, and that the biophysical properties of I_A_ and I_H_ are likely coregulated in mouse SNc DA neurons.

**SIGNIFICANCE STATEMENT:** Substantia nigra pars compacta (SNc) dopaminergic (DA) neurons are characterized by pacemaking activity, a broad action potential and biphasic post-inhibitory response. The A-type transient potassium current (I_A_) plays a central role in both pacemaking activity and post-inhibitory response. While it was thought so far that Kv4.3 ion channels were fully responsible for I_A_, using a Kv4.3^-/-^ transgenic mouse and antigen retrieval immunohistochemistry we demonstrate that Kv4.2 channels are also expressed in SNc DA neurons, although their contribution is significant in a minority of neurons only. Using electrophysiological recordings and computational modeling, we then demonstrate that I_A_ gating kinetics and its functional complementarity with the hyperpolarization-activated current are major determinants of both pacemaking activity and post-inhibitory response in SNc DA neurons.

## INTRODUCTION

While the expression of only two types of voltage-gated ion channels in the squid giant axon allowed Hodgkin and Huxley to dissect the biophysical processes underlying action potential (AP) genesis and conduction (Hodgkin and Huxley, 1952), most neuronal types express a multitude of ion channel subtypes underlying their electrical activity (Cembrowski et al., 2016; Fuzik et al., 2016; Tapia et al., 2018; Northcutt et al., 2019). In spontaneously active neurons, a variety of voltage- and calcium-gated ion channels are not only responsible for the AP, but also govern the subthreshold oscillations leading to AP firing, determine firing frequency and control its regularity (Atherton and Bevan, 2005; Swensen and Bean, 2005; Bean, 2007; Gantz et al., 2018). Substantia nigra pars compacta (SNc) dopaminergic (DA) neurons spontaneously generate a regular tonic pattern of activity, also known as “pacemaking” activity (Grace and Onn, 1989; Gantz et al., 2018). Over the past 40 years, many studies contributed to the identification of the specific ion channels involved in shaping pacemaking activity (Nedergaard and Greenfield, 1992; Liss et al., 2001; Seutin et al., 2001; Wolfart et al., 2001; Neuhoff et al., 2002; Liss et al., 2005; Chan et al., 2007; Puopolo et al., 2007; Guzman et al., 2009; Putzier et al., 2009; Ji et al., 2012; Gantz et al., 2018). In particular, several studies have suggested that the transient A-type potassium current (I_A_) plays an essential role in controlling pacemaking rate and post-inhibitory firing delay in these neurons (Liss et al., 2001; Putzier et al., 2008; Amendola et al., 2012; Tarfa et al., 2017). In addition, singlecell PCR, in situ hybridization and immunohistochemistry experiments suggested that the A-type current is carried exclusively by Kv4.3 ion channels (Serodio and Rudy, 1998; Liss et al., 2001; Ding et al., 2011; Dufour et al., 2014a; Tapia et al., 2018). Interestingly, several studies also suggested that the H-type current (I_H_, carried by HCN channels) displays strong functional interactions with I_A_, having for instance an opposite influence on post-inhibitory rebound delay (Amendola et al., 2012; Tarfa et al., 2017). The gating properties of these two currents were also shown to be co-regulated in rat SNc DA neurons (Amendola et al., 2012).

In the current study, we pursued the investigation of the role of Kv4 channels in the firing of SNc DA neurons. We first used electrophysiological recordings from wildtype (WT) and Kv4.3^-/-^ mice and computational modeling to determine the specific contribution of Kv4.3 ion channels and disentangle the respective influence of I_A_ conductance level, voltage sensitivity and gating kinetics on SNc DA neuron firing. As several studies have shown functional interactions between I_A_ and I_H_, we also investigated how modifications in I_A_ properties may alter I_H_ influence on firing, and vice-versa. Using computational modeling, we first found that spontaneous firing rate and post-inhibitory rebound are most sensitive to I_A_ maximal conductance and voltage-dependence. However, electrophysiological recordings obtained from WT and Kv4.3^-/-^ neurons suggested that pacemaking frequency and rebound delay in real neurons are mainly controlled by I_A_ gating kinetics. A very substantial reduction in I_A_ amplitude and inactivation rate was observed in most Kv4.3^-/-^ neurons, together with a strong increase and a strong reduction in pacemaking frequency and post-inhibitory rebound delay, respectively. Using antigen retrieval immunohistochemistry, we also showed that ~5% of the SNc DA neurons display a strong plasma membrane expression of Kv4.2 channels, which stabilizes electrical phenotype in the face of Kv4.3 deletion. Interestingly, no compensation of Kv4.3 loss by other ion currents was observed, although locomotion and motor learning were only mildly altered. This study thus provides new elements about the ion channels carrying I_A_ in SNc DA neurons, and demonstrates that I_A_ gating kinetics are critical in defining the physiological influence of this current on the firing of SNc DA neurons.

## MATERIAL AND METHODS

### Animals

Female and male P15-P80 WT (n=68 animals) and Kv4.3^-/-^ (n=40, Deltagen) mice from C57BL6/J genetic background were housed with free access to food and water in a temperature-controlled room (24°C) on a 12:12 h dark–light cycle (lights on at 07:00 h). All efforts were made to minimize the number of animals used and to maintain them in good general health, according to the European (Council Directive 86/609/EEC) and institutional guidelines for the care and use of laboratory animals (French National Research Council).

### Behavioral Experiments

Female and male WT (n=11) and Kv4.3^-/-^ (n=13) mice aged P56-P63 at the start of the behavioral testing were used to evaluate changes in motor function. As no statistical difference in behavior could be detected between males and females, data from both sexes were pooled and analyzed as a single sample.

#### Locomotor and exploratory activities

Actimetry was monitored in individual activity chambers (20 cm × 11.2 cm × 20.7 cm) housed within a sound-attenuating cubicle and under homogeneous illumination (Imetronic, Pessac, France). Each chamber was equipped with two pairs of infrared photobeams located 1.5 and 7.5cm above the floor level of the chamber. The number of back-and-forth movements (animals breaking the lower photobeams) as well as the number of vertical movements (animals breaking the upper photobeams) were recorded in 5-min bins over 90 min. Numbers of back-and-forth movements (locomotion) and vertical movements (rearing) are shown as mean ± SEM for each time bin over the whole period of recording time. Locomotion and rearing activities over the whole 90-min period were normalized and displayed as a mean percentage of control littermates.

#### Motor learning

Motor learning was evaluated on the accelerating rotarod (10cm diameter rod) test at a speed of 5 to 40 rotations per min (RPM) for 5 min. On the first day, mice were allowed to freely explore the non-rotating apparatus for 60s and subsequently trained to hold on the rotating rod (5 RPM) for at least two 60s trials, each trial being separated by a 10min break. Mice were allowed to recover for one hour before the first test. The testing phase consisted in 10 consecutive trials on the accelerating rod separated by 15-min breaks that allowed consolidation of performance. Results are shown as the average latency to fall off the rod (mean ± SEM) at each trial. A *performance index* was calculated for each individual and consisted in the average latency of the last 3 trials divided by the average latency of the first 3 trials multiplied by 100 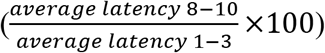.

### Electrophysiology

119 neurons from 39 WT mice and 109 neurons from 24 Kv4.3^-/-^ mice were recorded (current-clamp and voltage-clamp).

#### Acute midbrain slice preparation

Acute slices were prepared from P15-P25 animals of either sex. Mice were anesthetized with isoflurane (CSP) in an oxygenated chamber (TEM SEGA) before decapitation. After decapitation the brain was immersed briefly in oxygenated ice-cold low-calcium aCSF containing the following (in mM): 125 NaCl, 25 NaHCO3, 2.5 KCl, 1.25 NaH2PO4, 0.5 CaCl2, 4 MgCl2, and 25 D-glucose, pH 7.4, oxygenated with 95% O2/5% CO2 gas. The cortices were removed and then coronal midbrain slices (250 μm) were cut in ice-cold oxygenated low-calcium aCSF on a vibratome (Leica VT1200S and vibrating microtome 7000smz, Camden Instruments, UK). Following 20-30 min incubation in oxygenated low-calcium aCSF at 33°C, the acute slices were then incubated for a minimum of 30 min in oxygenated aCSF (containing in mM: 125 NaCl, 25 NaHCO3, 2.5 KCl, 1.25 NaH2PO4, 2 CaCl2, 2 MgCl2, and 25 glucose, pH 7.4, oxygenated with 95% O2/5% CO2 gas) at room temperature before electrophysiological recordings.

#### Drugs

Kynurenate (2 mM, Sigma-Aldrich) and picrotoxin (100 μM, Sigma-Aldrich) were used to block excitatory and inhibitory synaptic activity, respectively. AmmTX3 (1μM, Alomone) was used to block the transient potassium current (I_A_) carried by Kv4 channels. Drugs were bath applied via continuous perfusion in aCSF.

#### Electrophysiology recordings and analysis

All recordings (228 neurons from 63 mice) were performed on midbrain slices continuously superfused with oxygenated aCSF at 30–32°C. Picrotoxin and kynurenate were systematically added to the aCSF for all recordings to prevent contamination of the intrinsically generated activity by glutamatergic and GABAergic spontaneous synaptic activity. Patch pipettes (1.9–2.7 MOhm) were pulled from borosilicate glass (GC150TF-10, Harvard Apparatus) on a DMZ-Universal Puller (Zeitz Instruments) and filled with a patch solution containing the following (in mM): 20 KCl, 10 HEPES, 0.5 EGTA, 2 MgCl2, 0.4 Na-GTP, 2 Na2-ATP, 4 Mg-ATP,0.3 CaCl2, SUPERase RNase inhibitor (0.1 U/μl) and 115 K-gluconate, pH 7.4, 290-300 mOsm. For AmmTX3 experiments, patch pipettes (3.2-4.0 MOhm) were filled with a patch solution containing the following (in mM): 20 KCl, 10 HEPES, 0.5 EGTA, 2 MgCl2, 2 Na-ATP, and 120 K-gluconate, pH 7.4, 290-300 mOsm. Whole-cell recordings were made from SNc DA neurons visualized using infrared differential interference contrast videomicroscopy (QImaging Retiga camera; Olympus BX51WI microscope), and were identified based on their location, large soma size (>25μm), and electrophysiological profile (regular slow pacemaking activity, large spike half-width, large sag in response to hyperpolarizing current steps). For voltage-clamp experiments, only whole-cell recordings with an uncompensated series resistance <7 MOhm (compensated 85-90%) were included in the analysis. For current-clamp pharmacology experiments, higher series resistances were tolerated as long as the bridge compensation was properly adjusted to 100%. Liquid junction potential (−13.2 mV) and capacitive currents were compensated on-line. Recordings were acquired at 50kHz and were filtered with a low-pass filter (Bessel characteristic 2.8kHz cutoff). For current-clamp recordings, 1s hyperpolarizing current steps were injected to elicit a hyperpolarization-induced sag (due to I_H_ activation).

#### Current-clamp recordings and protocols

The spontaneous firing frequency was calculated from a minimum of 30 s of stable recording in cell-attached mode and from current-clamp recording (with no injected current) within the first 5 min after obtaining the whole-cell configuration. The coefficient of variation of the interspike interval (CVISI) was extracted from the same recording. Action potentials (APs) generated during this period of spontaneous activity were averaged and several parameters were extracted: AP threshold, AP amplitude, AP duration at half of its maximal height (AP half-width), AHP trough voltage, AHP latency. Hyperpolarizing current steps and depolarizing current steps were used to characterize the post-inhibitory rebound and the excitability properties. The gain start, gain end and spike frequency adaptation (SFA) index used to define excitability were calculated as described before (Dufour et al., 2014b).

#### Voltage-clamp recordings

The I_A_ current was elicited by a protocol consisting in a 500 ms prestep at −100 mV (to fully de-inactivate I_A_) followed by a 500 ms voltage step to −40 mV (to activate I_A_ without eliciting delayed rectifier potassium currents). The current generated by the same protocol using a prestep at −40mV (to fully inactivate I_A_) was subtracted to isolate I_A_. I_A_ properties (peak amplitude and total charge) were measured after subtracting the baseline at −40mV. The peak of the current elicited at −40mV was then plotted against the voltage of each corresponding prestep, and was fitted with a Boltzmann function to obtain I_A_ half-inactivation voltage (V_50_ I_A_) (see (Amendola et al., 2012)). The inactivation time constant (I_A_ tau) was extracted from a mono-exponential fit of the decay of the current. A two-step voltage-clamp protocol was used to determine the voltage-dependence of activation of I_H_ (V_50_ I_H_) and obtain the maximum I_H_ amplitude (see (Amendola et al., 2012) for details). For voltage-clamp recordings of delayed rectifier current (IKDR), tetrodotoxin (1 μM, Alomone), nickel (200 μM, Sigma-Aldrich) and cadmium (400 μM, Sigma-Aldrich) were also added to the aCSF. I_KDR_ was elicited by using a protocol consisting of a prestep at −30mV (to fully inactivate I_A_) followed by incremental depolarizing voltage steps up to +40mV.

#### Data acquisition

Data were acquired using an EPC 10 USB patch-clamp amplifier (HEKA) and the Patchmaster software acquisition interface (HEKA). Analysis was performed using FitMaster v2×73 (Heka).

### Immunohistochemistry

Adult (P21-P28) C57BL/6 WT mice (n=2) or Kv4.3^-/-^ littermates (n=2) of either sex were euthanized with ketamine-xylazine mix (100mg/Kg ketamine, 10mg/Kg xylazine), and transcardially perfused with PBS and ice-cold 4% paraformaldehyde in PBS. Brains were removed and post-fixed overnight (o/n) at 4°C in the same fixative solution. Coronal brain slices of 50 μm were obtained using a vibratome (vibrating microtome 7000smz, Camden Instruments, UK) and collected as floating sections. When indicated, antigen retrieval was performed by incubating the slices in sodium citrate (10mM, Sigma-Aldrich) during 30 min at 80°C (Jiao et al., 1999). Subsequently, slices were blocked for 1 h 30 min at room temperature (RT) in a solution containing 0.3% Triton X-100 (SIGMA) and 5% Normal Goat Serum (NGS, Vector Laboratories) in PBS. After blocking, sections were incubated with primary antibodies in a solution containing 0.3% Triton X-100 and 1% NGS in PBS (o/n; 4°C). The following primary antibodies were used in this study: Chicken anti-TH (1:1000; Abcam, ab76442, RRID:AB_1524535), Rabbit anti Kv4.3 (1:500, Alomone Labs, APC-017, RRID:AB_2040178), Mouse anti Kv4.2 (1:200, Neuromab, 75-361, clone L28/4, RRID:AB_2750684). After three washes (15 min/each) in PBS containing 0.3% Triton X-100, the floating sections were incubated with the following secondary antibodies: Alexa 488-Goat anti-mouse (1:200, Jackson ImmunoResearch), Alexa 488-Goat anti-rabbit (1:200, Jackson ImmunoResearch) and Alexa 594-Goat anti-chicken (1:200, Jackson ImmunoResearch) in a PBS solution containing 0.3% Triton X-100 and 1% NGS for 2 h at RT. Finally, sections were washed three times (15min/each), incubated with DAPI (1.5 μg/mL; Sigma-Aldrich) for 10 min, and mounted in Vectashield (Vector Laboratories). Sections were stored at 4°C, and images were acquired on a Zeiss LSM-780 confocal scanning microscope. All experiments involving WT and Kv4.3^-/-^ comparisons were performed in parallel applying the same acquisition settings to both genotypes. Images were processed and analyzed with ImageJ (NIH). Kv4.2-positive cells were visually identified in both genotypes on the basis of a perimembranous-like Kv4.2 staining and expressed as a percentage of the total number of TH+ cells. In order to compare the labeling pattern of Kv4.2 and Kv4.3, the line selection tool was used to trace 3μm-length lines perpendicular to the cell perimeter in individual optical sections. In each cell, 3 regions were analyzed, and 5 cells were used to calculate the average profile in each condition. Raw intensity values were collected, normalized (0-1 range) to the maximal value, and plotted as a function of distance (0 corresponding to Kv4 peak fluorescence signal, negative distances to extracellular space and positive distances to intracellular space; see **Figure 6B**). All the images shown are one single optical slice. For the sake of representation and to overcome differences regarding levels of TH expression in different DA neurons, different minimum and maximum display settings were applied in **Figure 6C** exclusively for the TH channel.

### Modeling

Simulations were performed using NEURON 7.5 software (Hines and Carnevale, 2001) as previously described (Moubarak et al., 2019). Realistic morphologies of 22 rat SNc DA neurons obtained previously were used to build multicompartment models (Moubarak et al., 2019). For each compartment, membrane voltage was obtained as the time integral of a first-order differential equation:

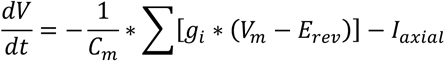

where V_m_ is the membrane potential, C_m_ the membrane capacitance, g_i_ are ionic conductances and E_rev_ their respective reversal potentials. The axial flow of current (I_axial_) between adjacent compartments is calculated by the NEURON simulation package (Hines and Carnevale, 2001). Cytoplasmic resistivity, specific membrane capacitance and specific membrane resistance were set to 150 Ohm.cm, 0.75 μF/cm^2^, and 100,000 Ohm.cm^2^, respectively, with E_rev_ for the leak conductance set at −50 mV. Six active conductances were included in the model: fast sodium (I_Na_), delayed rectifier potassium (I_KDR_), transient potassium (I_A_), L-type calcium (I_CaL_), hyperpolarization-activated (I_H_) and small conductance calcium-activated potassium (I_SK_) currents (Moubarak et al., 2019). Active conductances followed activation-inactivation Hodgkin-Huxley kinetics (**Table 1**). Parameters for I_A_, I_CaL_, I_SK_, I_Na_, I_KDR_ and I_H_ were based on our previous model and published values for SNc DA neurons (Gentet and Williams, 2007; Seutin and Engel, 2010; Amendola et al., 2012; Philippart et al., 2016; Moubarak et al., 2019). Intracellular calcium uptake was modeled as a simple decaying model according to Destexhe (Destexhe et al., 1993). Conductance values were set according to our own measurements or published values (see **Table 1**). Consistent with the literature (Kole and Stuart, 2008; Hu et al., 2009), g_Na_ and g_KDR_ were set to higher values in the axon initial segment (AIS) than in the rest of the neuron so that the action potential always initiated in the AIS. For sake of simplicity, activation and inactivation kinetics of I_A_ were voltage-independent but coupled to each other, such that activation rate was 50 times faster than inactivation rate.

**Table 1.**
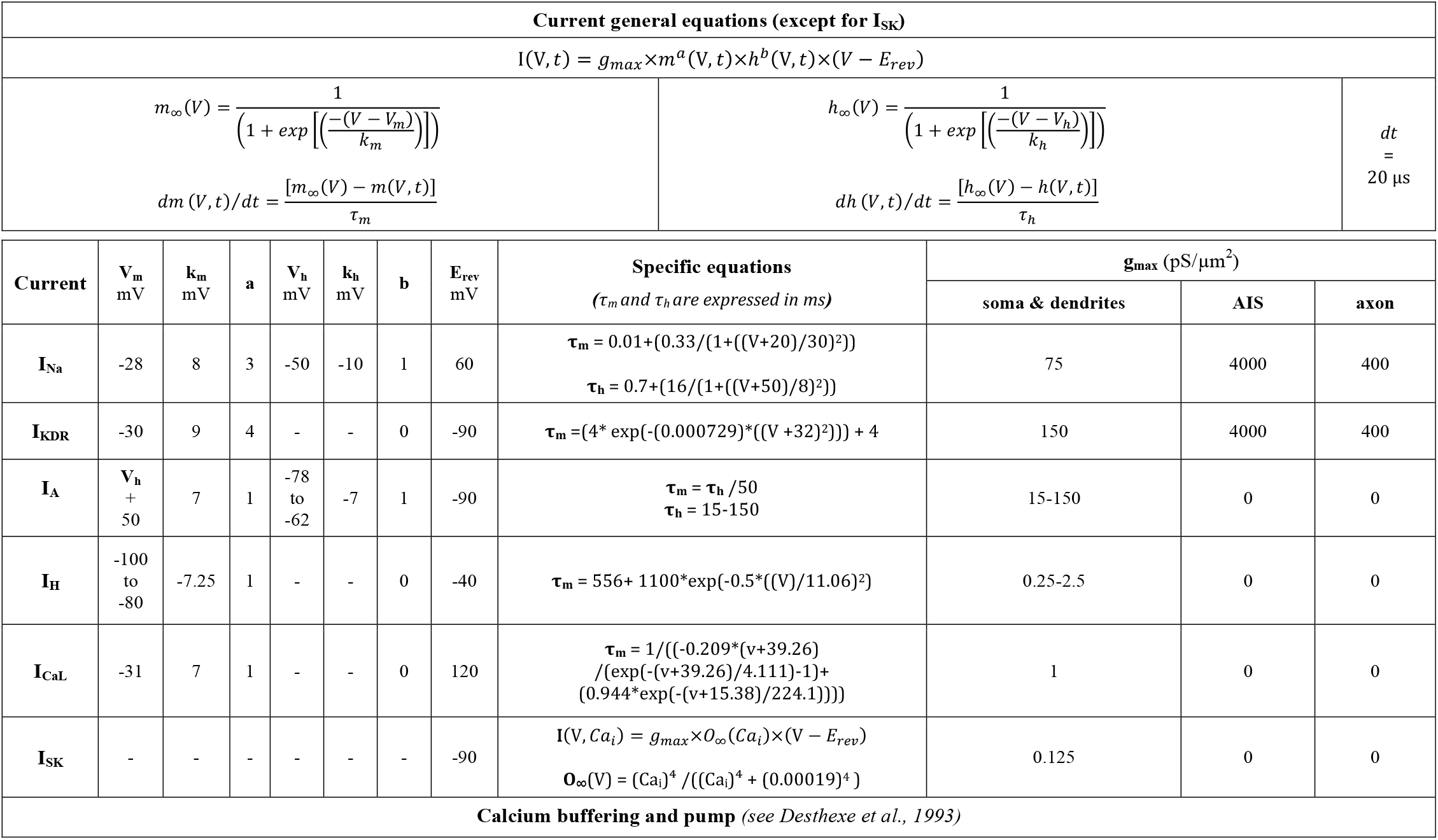
Equations governing the voltage dependence and kinetics of currents in the model.

In addition the inactivation and activation V_50s_ were also coupled (50 mV shift). As I_A_ and I_H_ voltage-dependences have been shown to be positively correlated in rat SNc DA neurons (Amendola et al., 2012), both values were forced to co-vary in the model according to the equation V_50_ inact. I_A_ = 0.814*(V_50_ act. I_H_)+3.36.

Initializing potential was set at −70 mV and pacemaking frequency was let to stabilize (4 spikes) before further analysis. Each simulation run had a duration of 8000 ms with a dt of 0.02 ms. Spatial discretization followed the “d_lambda rule” (Hines and Carnevale, 2001). All dendritic compartments and the axon-start compartment contained all currents whereas AIS and axon only contained fast sodium (I_Na_) and delayed rectifier potassium (I_KDR_) currents. To measure post-inhibitory rebound delay, current injection was performed by inserting a virtual electrode in the soma. A 1s pulse of current was injected into the model. Negative current amplitude was adjusted to achieve a peak hyperpolarization around −120 mV in each neuron and condition. Firing frequency, rebound delay and action potential property analyses were computed online by handmade routines directly written in NEURON hoc language (Hines and Carnevale, 2001). This model is derived from a previous model available at model DB database under the number 245427.

### Statistics

#### Behavior

For actimetry assessment, the numbers of horizontal (locomotion) and vertical (rearing) photobeam breaks were measured and compared between genotypes. Data are represented as line and scatter plots for the number of horizontal and vertical photobeam breaks per 5-min bin. The number of total movements was then normalized to the level of WT spontaneous locomotion. To assess the locomotor phenotype of Kv4.3^-/-^ mice, twoway repeated measures ANOVA tests with groups (WT/Kv4.3^-/-^) as the independent between-factor and time as the within-factor (training sessions or time-bins for locomotion) were performed. When the ANOVA was significant, multiple comparisons (Sidak post-hoc analyses) were used to evaluate differences between groups at different time points. P-values <0.05 were considered as statistically significant for all analyses. For motor learning, the average latency to fall off the accelerating rotarod was measured for each trial. Statistical difference in motor learning was assessed by comparing the *performance index*. Data are represented as line and scatter plots for the average latency to fall off the rod. Data are represented as mean ± SEM. After significant ANOVA, post-hoc unpaired t-tests between Kv4.3^-/-^ mice and their respective WT littermates were performed for the actimetry and rotarod behavioral tests. Statistical analyses were performed using Prism6 (GraphPad Software Inc., La Jolla, USA).

#### Electrophysiology and immunohistochemistry

The univariate statistical analysis of electrophysiological data, performed according to the distribution properties of the data using a Shapiro-Wilk normality test, included paired t-test or Wilcoxon signed rank test; t-test or Mann-Whitney Wilcoxon test with p < 0.05 considered to be statistically significant (R environment (R Core Team 2016, (https://www.r-project.org/)), SigmaPlot 11.0, GraphPad Prism 6). In most figures, data are represented as scatter plots or box and whisker plots, with all individual points appearing on the graphs and dotted lines indicating the distribution of data (violin plots). For pharmacological experiments, data are represented as mean ± SEM (scatter or bar plots). Correlation, linear regression and multiple linear regression analysis (**Figure 1**, **9**) were performed in R. For every pair of variables, correlation parameters, rho (Spearman correlation factor) or r (Pearson correlation factor), were selected after performing a Shapiro-Wilk normality test on the linear regression residuals and p values were corrected for multiple comparisons by a false discovery rate adjustment (stats package; (Benjamini and Hochberg, 1995)). For multiple linear regression, variables (extracellular ISI, rebound delay, I_A_ tau, I_H_ amplitude and I_A_ amplitude) were first log transformed, and then dependent variables were standardized (subtracting the mean and dividing by the SD). A selection of the best subsets of dependent variables for each model size (1 to 4 for the model and 1 to 5 for real data) was first performed (leaps package) according to several criteria (adjusted R^2^, AIC, BIC). The best model was then selected by comparing the prediction error of each model after performing a repeated (20 times) 10-fold crossvalidation on test data (caret package). The best linear model, corresponding to the minimum cross-validation error (i.e a model with the best predictive power) was then obtained. Multicolinearity was assessed by computing a score called the variance inflation factor (VIF package) and VIF was < 1.5 for all variables retained in the different models. For the immunohistochemistry experiments, a Fisher exact test was used to compare the proportions of Kv4.2-positive cells among TH-positive cells in WT and Kv4.3^-/-^ mice. Figures were prepared using R, SigmaPlot 11.0, GraphPad Prism 6 and Adobe Illustrator CS5/CS6.

**Figure 1.**
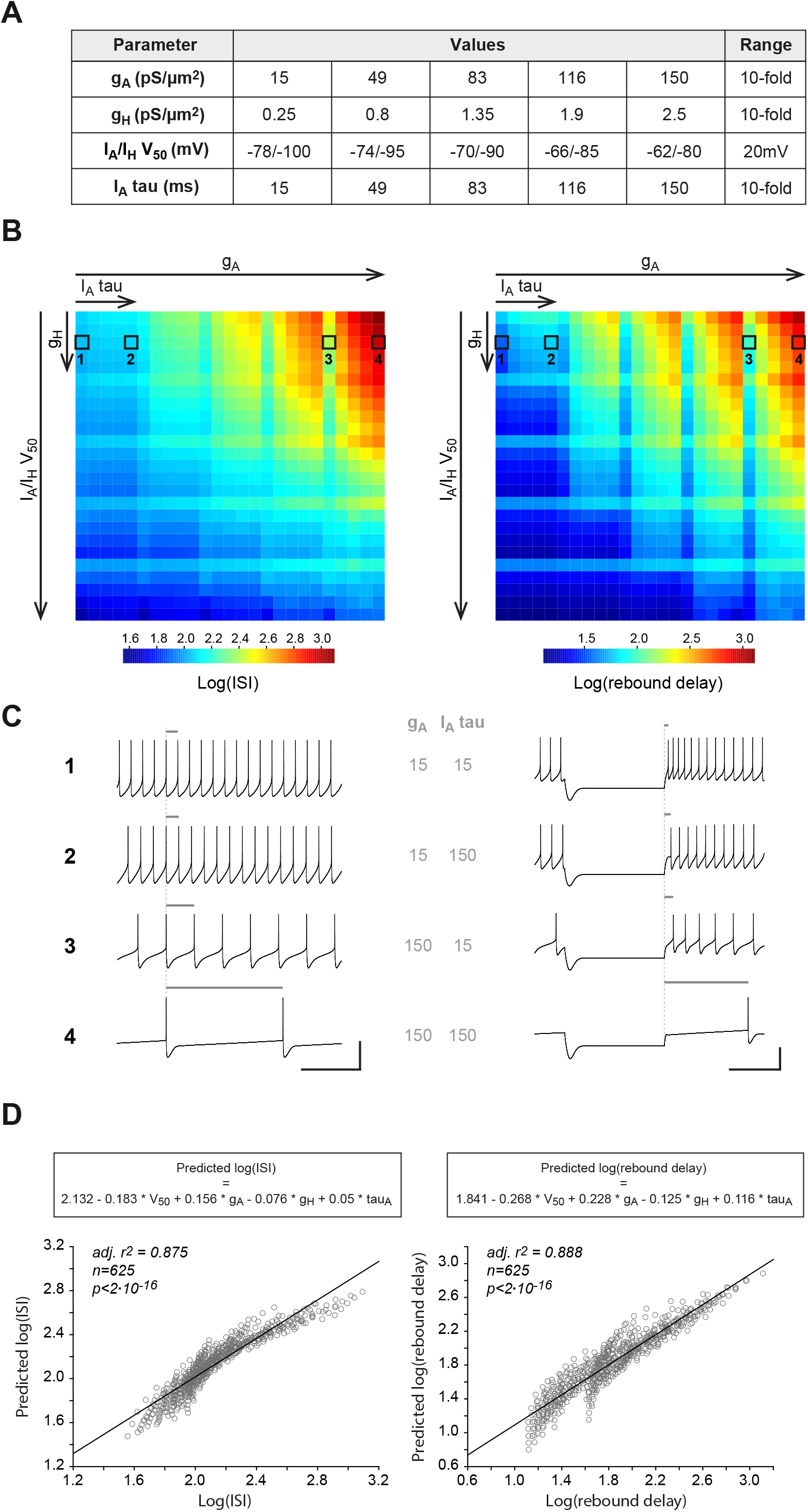
Modeling the effect of the biophysical properties of I_A_ and I_H_ on spontaneous activity and post-inhibitory rebound. **A**, table presenting the 5 values tested for each biophysical parameter (g_A_, g_H_, I_A_/I_H_ V_50_ and I_A_ inactivation tau) in the multi-compartment model. Each property was varied independently, leading to 625 (5^4^) versions of the model. **B**, dimensionally stacked heatmaps showing the variation in ISI (displayed as log(ISI), left) and in post-inhibitory rebound delay (displayed as log(rebound delay), right) as a function of g_A_ (X axis, big scale), I_A_/I_H_ V_50_ (Y axis, big scale), I_A_ tau (X axis, small scale) and g_H_ (Y axis, small scale). **C**, example traces of spontaneous pacemaking (left) and post-inhibitory rebound (right) obtained for the minimal and maximal values of g_A_ and I_A_ tau (corresponding to the 4 squares surrounded in the panel **B** heatmaps). The gray horizontal bars above the traces help visualize the change in ISI (left) or rebound delay (right) as a function of changes in g_A_ and I_A_ tau. **D**, multiple linear regression reveals the relative contribution of each biophysical property to ISI (left) or rebound delay variation (right). The scatter plots show the relationship between the measured values of log(ISI) and log(rebound delay) and the values predicted using a linear combination of the 4 biophysical variables listed in panel **A** (the corresponding equations are shown above each graph). Scale bars: **C**, horizontal 500ms, vertical 50mV.

## RESULTS

### Influence of I_A_ biophysical properties on SNc DA neuron firing

Several studies have investigated the role of I_A_ in SNc DA neurons (Liss et al., 2001; Putzier et al., 2008; Amendola et al., 2012; Tarfa et al., 2017). One of the first studies investigating the molecular substrate of this current suggested that the cell-to-cell variation in I_A_ charge density, mainly due to variation in Kv4.3 expression level, had a strong influence on pacemaking frequency (Liss et al., 2001). Two other studies investigated more specifically the influence of I_A_ on post-inhibitory firing and concluded that both its voltage-dependence and inactivation rate had a major influence on rebound firing delay (Amendola et al., 2012; Tarfa et al., 2017). In addition, these latter studies highlighted the complementary influence of I_H_ on rebound firing. For instance, I_A_ and I_H_ voltage-dependences were found to co-vary in rat SNc DA neurons, this co-variation defining the dynamic range of post-inhibitory rebound delay (Amendola et al., 2012). In order to determine which of the properties of I_A_ plays the most critical role in the control of spontaneous firing rate and post-inhibitory firing, we used a realistic multicompartment Hodgkin-Huxley model of rat SNc DA neurons (Moubarak et al., 2019). Based on measurements obtained by different groups (Liss et al., 2001; Gentet and Williams, 2007; Amendola et al., 2012; Tarfa et al., 2017), each of the biophysical properties of I_A_ (maximal conductance g_A_, voltage-dependence I_A_ V_50_, inactivation rate I_A_ tau) and I_H_ maximal conductance (g_H_) were varied over a 10-fold range (20mV range for the voltage-dependence) using 5 equi-distributed values (**Figure 1A**). As I_A_ and I_H_ voltage-dependences have been shown to be positively correlated in rat SNc DA neurons (Amendola et al., 2012), both values were forced to co-vary in the model. Using a sample of 22 realistic models and 5 independently-varying values for each biophysical property, a database of 13750 models (22*5^4^) was generated (**Figure 1**). The average interspike interval (ISI) during spontaneous activity and the post-inhibitory firing delay in response to a hyperpolarizing pulse (rebound delay) were measured for each model and their average values were calculated for each combination of values of the 4 biophysical parameters (n=625). Dimensional stacking (Taylor et al., 2006) was then used to represent the influence of the 4 biophysical parameters on these electrophysiological features in 2-dimensional heatmaps, allowing us to visually determine which parameters were most critical in controlling ISI and rebound delay (**Figure 1B, C**): g_A_ and I_A_/I_H_ V_50_ strongly modulated both ISI and rebound delay while I_A_ tau and g_H_ had a weaker influence on these firing properties. To quantify the contribution of each biophysical parameter to ISI and rebound delay variations, we then standardized the values of each parameter (by subtracting the mean and dividing by the standard deviation) and used multiple linear regression against ISI or rebound delay (**Figure 1D**). Consistent with the visualization provided in **Figure 1B**, this sensitivity analysis revealed that the influence of g_A_ and I_A_/I_H_ V_50_ on ISI and rebound delay was 2-3 times stronger than that of g_H_ and I_A_ tau. While these simulations provide important insights into the potential influence of specific biophysical properties of I_A_ (and I_H_) on SNc DA neuron firing, we then sought to determine the precise role of I_A_ in these neurons ex silico by using a transgenic mouse lacking the main type of Kv4 channel (Kv4.3) thought to underlie the A-type current in midbrain DA neurons (Serodio and Rudy, 1998; Liss et al., 2001; Amendola et al., 2012; Dufour et al., 2014a).

### Phenotypic evaluation of the Kv4.3^-^ transgenic mouse

The expression of Kv4.3 is particularly high in midbrain DA neurons (Serodio et al., 1996; Serodio and Rudy, 1998; Liss et al., 2001; Dufour et al., 2014a). Consistent with these studies, we observed a high level of expression of Kv4.3 in the SNc of WT mice, with the expected membrane profile of immunostaining (**Figure 2A**). The immunolabeling was completely lost in the SNc of Kv4.3^-/-^ mice (**Figure 2A**). Although the Kv4.3 knock-out mouse used here has already been studied previously (Niwa et al., 2008; Carrasquillo et al., 2012), its behavioral phenotype has not been thoroughly evaluated. As Kv4.3 is strongly expressed in the SNc, we sought to determine whether Kv4.3 loss could affect SNc-related behaviors, such as locomotion and motor learning (**Figure 2B-D**). As illustrated in **Figure 2B-C**, locomotor activity (horizontal and vertical movements) showed a steady decrease over time, which was similar between Kv4.3^-/-^ mice and their WT littermates, indicating habituation to the actimetry chamber. Consistently, the average horizontal and vertical movements recorded over 90 min were not significantly different between Kv4.3^-/-^ mice and WT littermates (**Figure 2C**). Motor learning abilities were then assessed using the rotarod test for ten consecutive trials (**Figure 2D**). While WT mice improved their performance across trials, as shown by the increase in the latency-to-fall, Kv4.3^-/-^ mice showed a mild impairment in motor learning, as indicated by their significantly lower performance index in this motor task (176.9 ± 21.4, n=11 *vs* 111.2 ± 8.3, n=13, p=0.006, unpaired t-test, **Figure 2D**). Overall, the Kv4.3^-/-^ mice displayed moderate behavioral alterations, suggesting that basal ganglia-related motor functions are at least partially robust to the complete loss of Kv4.3.

**Figure 2.**
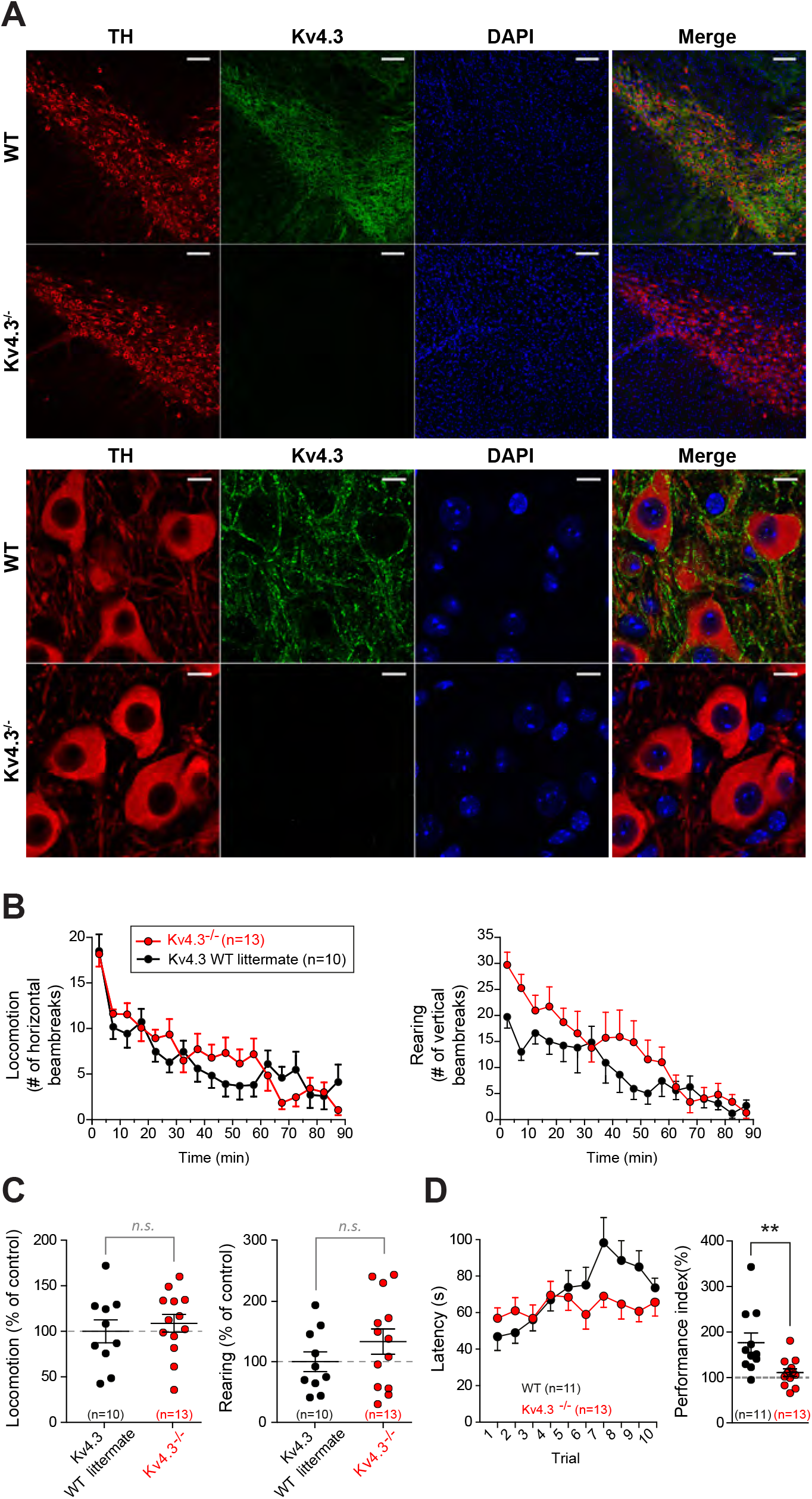
Validation and overall robustness of the Kv4.3^-/-^ transgenic mouse. **A**, Kv4.3 immunostaining in the SNc confirms that Kv4.3 expression is lost in Kv4.3^-/-^ mice. Top, low-magnification pictures of the SNc showing (from left to right) TH (red), Kv4.3 (green) and DAPI (blue) stainings in WT (top row) and Kv4.3^-/-^ mice (bottom row). Bottom, high-magnification pictures of SNc DA neurons showing (from left to right) TH (red), Kv4.3 (green) and DAPI (blue) stainings in WT (top row) and Kv4.3^-/-^ mice (bottom row). **B**, locomotion measured in actimetry chambers. Left, line and scatter plot showing the mean number of horizontal movements per 5-min bin (± SEM) in Kv4.3^-/-^ mice (red) compared to their WT littermates (black). Right, line and scatter plot showing the mean number of rearing events per 5-min bin in Kv4.3^-/-^ mice (red) compared to their WT littermates (black). **C**, scatter plots showing the total number of horizontal movements (left) or rearing events (right) per 90-min session in Kv4.3^-/-^ mice normalized to WT littermate average. **D**, changes in motor learning measured using a rotarod assay. The latency to falling off the rotating rod (with increasing rotating speed) was assessed over 10 consecutive trials (left). The performance index ((average latency 8-10)/(average latency 1-3)*100, right) was used to evaluate the learning ability of Kv4.3^-/-^ mice (red) compared to their WT littermates (black). ** p<0.01. Scale bars: **A**, low-magnification pictures 100μm, high-magnification pictures 10μm.

### Electrophysiological phenotype of Kv4.3^-/-^ SNc DA neurons

Following the approach already used in a previous study (Dufour et al., 2014b), we performed an exhaustive current-clamp characterization of the firing properties of SNc DA neurons to determine changes in phenotype associated with Kv4.3 deletion (**Table 2**). Passive properties, spontaneous activity, post-inhibitory rebound, action potential shape and excitability were assessed by measuring 16 different electrophysiological parameters in 75-101 WT and 66-77 Kv4.3^-/-^ neurons. The first obvious electrophysiological change observed was that spontaneous activity (extracellularly recorded in cell-attached mode) was dramatically modified in Kv4.3^-/-^ SNc DA neurons (**Figure 3A, B**). Spontaneous firing rate was increased by ~2-fold in Kv4.3^-/-^ mice, as demonstrated by the significant decrease in interspike interval (ISI, **Figure 3A, B**; **Table 2**). Pacemaking regularity, measured by the coefficient of variation of the ISI (CV_ISI_) was also significantly different in Kv4.3^-/-^ mice, although CV_ISI_ values were very low (<20%) in both WT and Kv4.3^-/-^ mice, indicating a highly regular tonic activity. Consistent with previous studies demonstrating the critical role of Kv4 channels in post-inhibitory rebound in SNc DA neurons (Amendola et al., 2012; Tarfa et al., 2017), post-inhibitory rebound delay was also dramatically decreased in Kv4.3^-/-^ mice (**Figure 3C, D**; **Table 2**). However, the I_H_- mediated voltage sag observed during prolonged hyperpolarization was not modified (**Figure 3C, D**; **Table 2**). In spite of the demonstrated role of Kv4 channels in controlling spike shape in other cell types (Nerbonne et al., 2008; Carrasquillo et al., 2012), most action potential parameters were unchanged in Kv4.3^-/-^ mice, except for action potential half-width, which was slightly larger (**Figure 4A**; **Table 2**). We also analyzed neuronal excitability by measuring the responses of the neurons to increasing depolarizing current steps (**Figure 4B**). As SNc DA neurons display firing frequency adaptation in response to prolonged depolarization (Vandecasteele et al., 2011; Dufour et al., 2014b), gain was quantified on the first (gain start) and last ISI (gain end), and a spike frequency adaptation index could be computed (SFA index = gain start/gain end). Consistent with previous observations made on cortical neurons (Carrasquillo et al., 2012), excitability was slightly increased in Kv4.3^-/-^ neurons, although this change only affected the initial response of neurons (gain start) to current injection (**Figure 4B**; **Table 2**). Consistently, the frequency of the response of Kv4.3^-/-^ neurons to a 100pA step was also found to be significantly higher (**Figure 4B**, **Table 2**).

**Figure 3.**
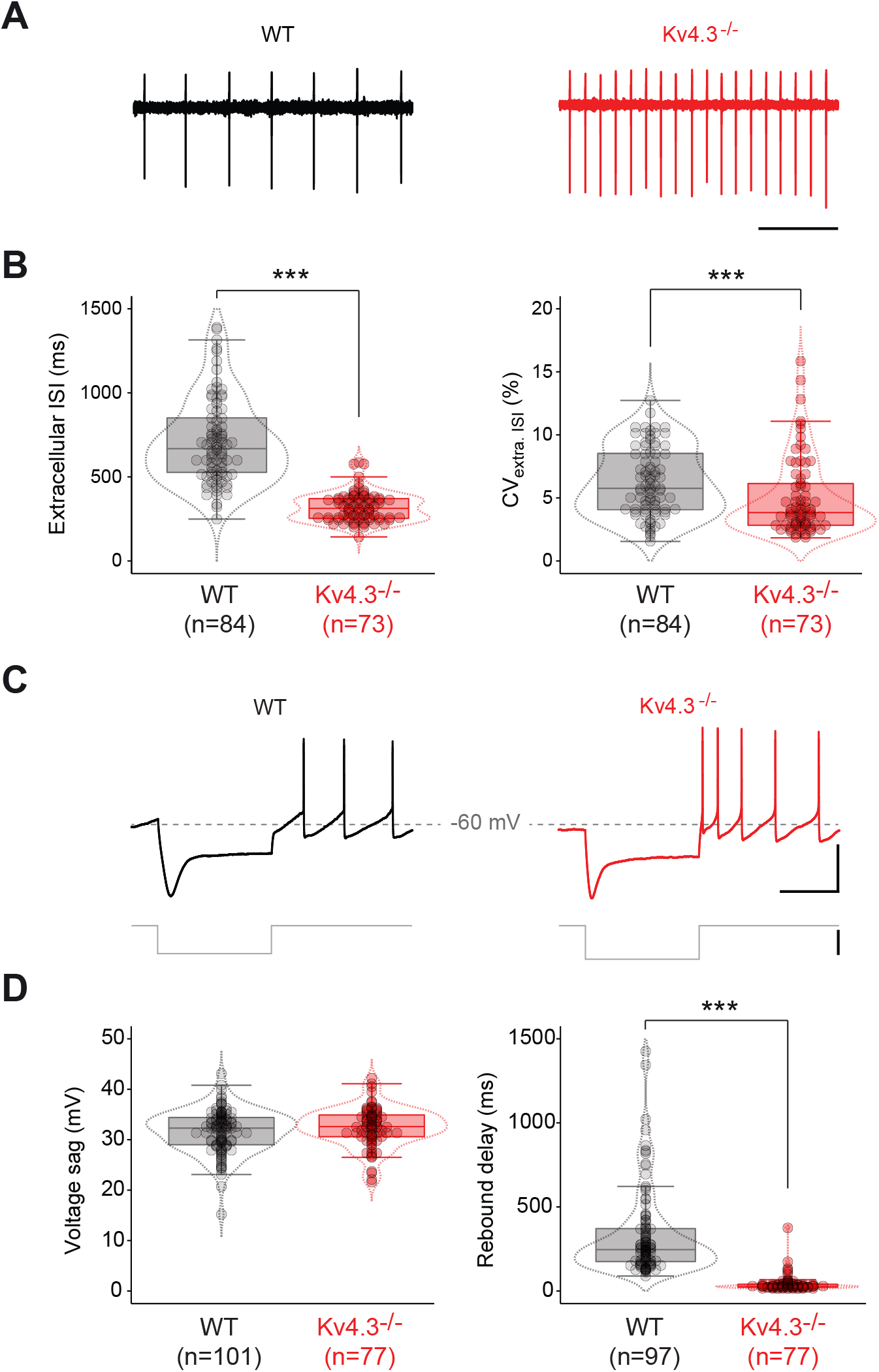
Spontaneous activity and post-inhibitory rebound are profoundly altered in Kv4.3^-/-^ SNc DA neurons. **A**, representative cell-attached recordings showing the spontaneous pattern of activity in WT (black trace, left) and Kv4.3^-/-^ SNc DA neurons (red trace, right). **B**, box and whisker plots showing the distribution of values for extracellularly recorded spontaneous ISI (Extracellular ISI, left) and ISI coefficient of variation (CV_extra_. _ISI_, right) in WT and Kv4.3^-/-^ SNc DA neurons. **C**, representative current-clamp recordings showing the voltage response of SNc DA neurons to a current step (gray trace) hyperpolarizing membrane voltage to ~ ‒120mV in WT (black trace, left) and Kv4.3^-/-^ mice (red trace, right). The recordings come from the same neurons as the cell-attached recordings presented in **A**. **D**, box and whisker plots showing the distribution of values for voltage sag amplitude (left) and post-inhibitory rebound delay (right) in WT and Kv4.3^-/-^ SNc DA neurons. *** p<0.001. Dotted lines in the box and whisker plots indicate the distribution of data (violin plots). Scale bars: **A**, horizontal 1s; **C**, top, horizontal 500ms, vertical 40mV; bottom, vertical 100pA; horizontal dotted lines indicate -60mV.

**Figure 4.**
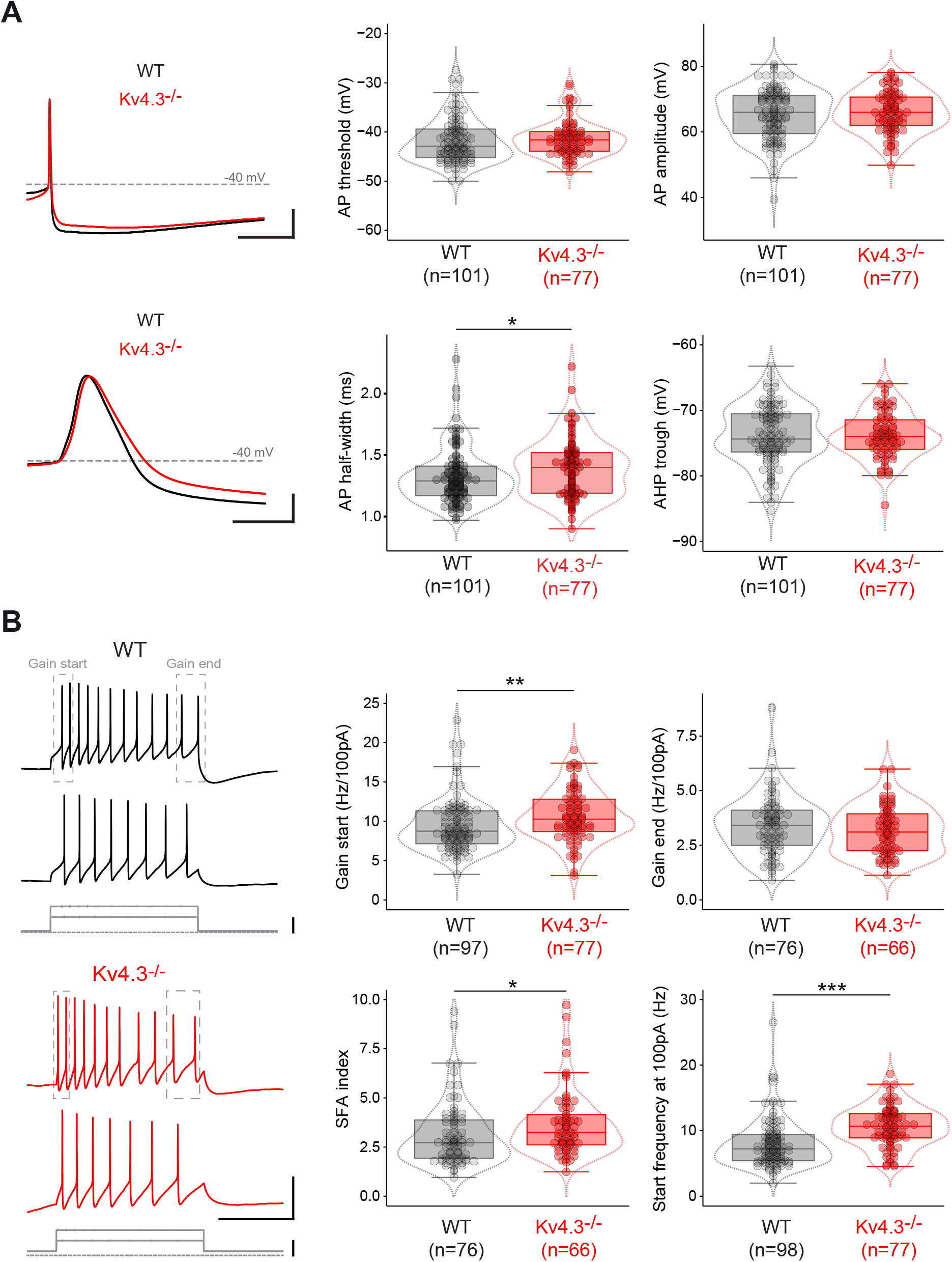
Action potential and excitability properties in Kv4.3^-/-^ SNc DA neurons. **A**, left, current-clamp recordings showing the shape of the action potential in the WT (black traces) and Kv4.3^-/-^ mice (red traces) on a slow (top) and fast time-scale (bottom). Right, box and whisker plots showing the distribution of values for action potential threshold (top left), amplitude (top right), half-width (AP half-width, bottom left) and afterhyperpolarization trough (AHP trough, bottom right) in WT and Kv4.3^-/-^ SNc DA neurons. **B**, left, current-clamp recordings showing the voltage response of SNc DA neurons to 100 and 200pA depolarizing current steps (gray traces) in WT (top, black traces) and Kv4.3^-/-^ mice (bottom, red traces). Gray dotted rectangles indicate the interspike interval used to calculate the gain start and gain end on each train of action potentials. Right, box and whisker plots showing the distribution of values for gain start (top left), gain end (top right), spike frequency adaptation (SFA) index (bottom left) and start frequency at 100pA (bottom right) in WT and Kv4.3^-/-^ SNc DA neurons. * p<0.05, ** p<0.01, *** p<0.001. Dotted lines in the box and whisker plots indicate the distribution of data (violin plots). Scale bars: **A**, top, horizontal 50ms, vertical 20mV; **A**, bottom, horizontal 2ms, vertical 20mV, horizontal dotted lines indicate −40mV; **B**, voltage, horizontal 500ms, vertical 40mV; current, vertical 100pA.

**Table 2.** Statistical analysis of electrophysiological parameters in wild-type and Kv4.3^-/-^ SNc DA neurons. The values for 16 electrophysiological parameters measured under current-clamp (corresponding to passive properties, spontaneous activity, post-inhibitory rebound, action potential and excitability) and 8 electrophysiological parameters measured under voltage-clamp (corresponding to I_A_ and I_H_ properties) are presented for WT and Kv4.3^-/-^ SNc DA neurons. Mean and SD (black text) are reported for normally-distributed data, while Median and interquartile range (IQR) are reported otherwise (gray text). Accordingly, statistical differences between WT and Kv4.3^-/-^ neurons were tested using a t-test or a Mann Whitney test, depending on the normality of the data. Asterisks indicate statistically significant differences (* p<0.05, ** p<0.01, *** p<0.001).

| Parameter | WT |  |  | Kv4.3<sup>-/-</sup> |  |  | p-value<br>(t-test)<br>(Mann Whitney) |
| --- | --- | --- | --- | --- | --- | --- | --- |
|  | Mean/Median | SD/IQR | n | Mean/Median | SD/IQR | n |  |
| **Passive properties** |  |  |  |  |  |  |  |
| R<sub>m</sub> (Mohm) | 386.00 | 202.00 | 97 | 422.00 | 147.00 | 72 | 0.2351 |
| **Pacemaking** |  |  |  |  |  |  |  |
| ISI (ms) | 669.50 | 334.50 | 84 | 304.00 | 118.00 | 73 | 2.2·10<sup>-16</sup> *** |
| CV<sub>ISI</sub> (%) | 5.76 | 4.45 | 84 | 3.84 | 3.31 | 73 | 4·10<sup>-4</sup> *** |
| **Post-inhibitory rebound** |  |  |  |  |  |  |  |
| Voltage sag (mV) | 31.70 | 4.29 | 101 | 32.50 | 3.95 | 77 | 0.2082 |
| Rebound delay (ms) | 256.00 | 199.00 | 97 | 29.00 | 18.00 | 77 | 2.2·10<sup>-16</sup> *** |
| **Action potential** |  |  |  |  |  |  |  |
| AP Threshold (mV) | -42.90 | 5.80 | 101 | -41.60 | 4.00 | 77 | 0.15 |
| AP amplitude (mV) | 65.29 | 7.84 | 101 | 66.01 | 6.27 | 77 | 0.5024 |
| AP half-width (ms) | 1.29 | 0.24 | 101 | 1.40 | 0.33 | 77 | 0.038 * |
| AP rise slope (mV/ms) | 86.47 | 24.95 | 101 | 87.89 | 24.61 | 77 | 0.7049 |
| AP decay slope (mV/ms) | -47.71 | 8.73 | 101 | -45.33 | 8.89 | 77 | 0.0756 |
| AHP trough voltage (mV) | -73.84 | 4.58 | 101 | -73.83 | 3.58 | 77 | 0.9892 |
| AHP latency (ms) | 55.56 | 38.28 | 101 | 53.76 | 28.68 | 77 | 0.84 |
| **Excitability** |  |  |  |  |  |  |  |
| Start frequency at 100 pA (Hz) | 7.23 | 3.98 | 98 | 10.68 | 3.73 | 77 | 4·10<sup>-8</sup> *** |
| Gain start (Hz/100pA) | 8.75 | 4.18 | 97 | 10.27 | 4.11 | 77 | 1.7·10<sup>-3</sup> ** |
| Gain end (Hz/100pA) | 3.41 | 1.61 | 76 | 3.11 | 1.69 | 66 | 0.37 |
| SFA index | 2.73 | 1.94 | 76 | 3.24 | 1.54 | 66 | 0.012 * |
| **I<sub>A</sub>** |  |  |  |  |  |  |  |
| V<sub>50</sub> inactivation (mV) | -68.91 | 5.10 | 87 | -73.31 | 3.84 | 39 | 4.3·10<sup>-6</sup> *** |
| Inactivation tau (ms) | 30.90 | 27.15 | 83 | 12.90 | 6.72 | 42 | 7.1·10<sup>-12</sup> *** |
| Amplitude (nA) | 2.14 | 2.79 | 87 | 0.93 | 0.64 | 39 | 1.8·10<sup>-12</sup> *** |
| Charge (pAs) | 152.80 | 177.80 | 83 | 19.40 | 22.40 | 37 | 6.4·10<sup>-15</sup> *** |
| Charge density (pAs/pF) | 1.17 | 1.00 | 82 | 0.18 | 0.20 | 36 | 6.4·10<sup>-13</sup> *** |
| **I<sub>H</sub>** |  |  |  |  |  |  |  |
| V<sub>50</sub> activation (mV) | -88.41 | 4.24 | 75 | -88.42 | 3.81 | 55 | 0.9886 |
| Amplitude (pA) | 453.00 | 212.50 | 75 | 575.30 | 275.80 | 57 | 6·10<sup>-4</sup> *** |
| Amplitude density (pA/pF) | 3.49 | 1.93 | 74 | 5.76 | 2.36 | 54 | 7.6·10<sup>-11</sup> *** |

### Voltage-clamp characterization of I_A_ in Kv4.3^-^ SNc DA neurons

We then directly investigated changes in the properties of I_A_ by performing voltage-clamp recordings in WT and Kv4.3^-/-^ SNc DA neurons (**Figure 5**). A dramatic decrease in I_A_ amplitude was observed in Kv4.3^-/-^ mice (**Figure 5A, B**; **Table 2**). However, a small transient residual current with I_A_-like properties (voltage-dependent inactivation) was still present in all Kv4.3^-/-^ recordings (**Figure 5A**). Most interestingly, this residual current was completely blocked by the Kv4-specific toxin AmmTX3 (n=4, no measurable residual current after toxin application; **Figure 5A**), suggesting that a Kv4 subunit other than Kv4.3 is expressed at a low level in Kv4.3^-/-^ SNc DA neurons. As the impact of I_A_ on firing strongly depends on its gating kinetics (Amendola et al., 2012; Tarfa et al., 2017), we then measured its time constant of inactivation (I_A_ tau) and calculated the overall charge carried by the current (**Figure 5C-E**). Both parameters were strongly decreased in Kv4.3^-/-^ SNc DA neurons (**Figure 5C, D**; **Table 2**), although a minority of cells (n=5/42) displayed values similar to the WT measurements for both of these parameters. Plotting I_A_ charge versus I_A_ tau revealed the clear separation of values between the Kv4.3^-/-^ and WT measurements, except for the 5 cells identified before (**Figure 5E**). Based on these voltage-clamp data, it appears that, although the Kv4.3 subunit by far predominates in WT SNc DA neurons, another Kv4 subunit is also expressed, at least in the Kv4.3^-/-^ neurons. Although in most cases, the expression level of this unidentified subunit is too low to compensate for the loss of Kv4.3, it generates an A-type current that provides a minority of Kv4.3^-/-^ SNc DA neurons (5/42 = 12%) with a “wild-type” voltage-clamp phenotype.

**Figure 5.**
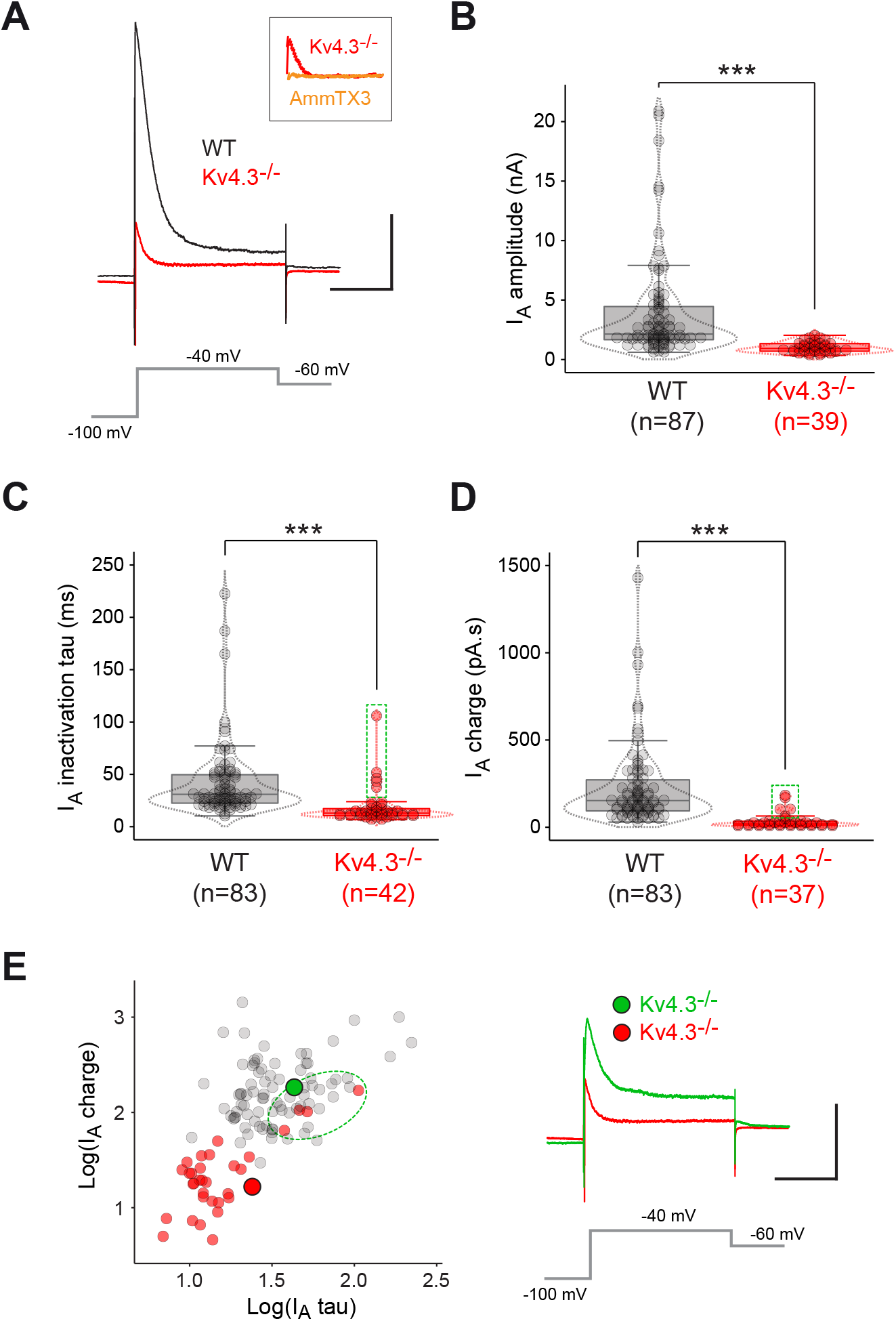
Voltage-clamp analysis of I_A_ in wild-type and Kv4.3^-/-^ SNc DA neurons. **A**, voltage-clamp traces showing representative I_A_ recordings obtained from a WT (black trace) and a Kv4.3^-/-^ SNc DA neuron (red trace) in response to a voltage step to −40mV (gray trace). The small residual current present in the Kv4.3^-/-^ mice is blocked by AmmTX3 (inset, orange trace). **B**, box and whisker plot showing the distribution of values for I_A_ amplitude in WT and Kv4.3^-/-^ SNc DA neurons. **C**, box and whisker plot showing the distribution of values for I_A_ time constant of inactivation (I_A_ tau) in WT and Kv4.3^-/-^ SNc DA neurons. The green dotted rectangle highlights 5 Kv4.3^-/-^ outliers displaying unusually large values for I_A_ tau. **D**, box and whisker plot showing the distribution of values for I_A_ charge in WT and Kv4.3^-/-^ SNc DA neurons. The green dotted rectangle highlights 5 Kv4.3^-/-^ outliers displaying unusually large values for I_A_ charge (same cells as in **C**). **E**, left, scatter plot showing the relationship between I_A_ tau and charge in WT (gray dots) and Kv4.3^-/-^ SNc DA neurons (red dots). Please note that 5 of the Kv4.3^-/-^ measurements lie in the WT region of space (green dotted ellipse). Right, voltage-clamp traces showing one example of the atypical I_A_ recording (green trace, corresponding to the large green circle in the scatter plot) encountered in one of the 5 Kv4.3^-/-^ outliers highlighted in panels **C** and **D**, compared to the typical recording obtained in Kv4.3^-/-^ neurons (red trace, same as in panel **A**, corresponding to the large red circle in the scatter plot). *** p<0.001. Dotted lines in the box and whisker plots indicate the distribution of data (violin plots). Scale bars in **A** and **E**: horizontal 200ms, vertical 1nA.

### Using antigen retrieval to reveal the expression of Kv4.2 channels by SNc DA neurons

Several studies have investigated the expression of A-type Kv channels in SNc DA neurons, using in situ hybridization (Serodio et al., 1996; Serodio and Rudy, 1998), single-cell quantitative PCR (Liss et al., 2001; Ding et al., 2011; Tapia et al., 2018) and immunohistochemistry (Liss et al., 2001; Dufour et al., 2014a). While a high level of expression for Kv4.3 (at both the mRNA and protein level) (Liss et al., 2001; Dufour et al., 2014a; Tapia et al., 2018) and the absence of Kv4.1 (Serodio and Rudy, 1998; Liss et al., 2001; Ding et al., 2011) were consistently reported, the presence of Kv4.2 is subject to more debate. In particular, Kv4.2 mRNA has been detected in several studies (Ding et al., 2011; Tapia et al., 2018), although the protein was not detected by classical immunohistochemistry (Liss et al., 2001; Dufour et al., 2014a). Interestingly, it has been shown that, depending on the brain region and the subcellular location of the ion channel of interest, an antigen retrieval (AR) procedure may be required to uncover potassium channel antigen epitopes before performing immunolabeling (Lorincz and Nusser, 2008). In order to test the efficiency of AR on Kv4.2 staining, we first performed Kv4.2 immunolabeling with or without AR on the CA1 region of the hippocampus where this ion channel is highly expressed (Serodio and Rudy, 1998). As can be seen in **Figure 6A**, Kv4.2 immunostaining was greatly improved by AR, revealing a strong perisomatic and dendritic staining of pyramidal cells. We therefore performed Kv4.2 immunostaining with and without AR on WT midbrain slices (**Figure 6B**). Similar to what we observed in the hippocampus, Kv4.2 immunostaining in the SNc was greatly improved by AR, although only a minority of DA neurons (TH-positive) displayed a clear perisomatic Kv4.2 signal compatible with membrane expression of the channel. In fact, the AR Kv4.2 staining profile of Kv4.2-positive cells was very similar to the membrane staining profile observed for Kv4.3 (**Figure 6B**). This distinctive staining profile was then used to quantify the percentage of Kv4.2-positive SNc DA neurons in both WT and Kv4.3^-/-^ mice (**Figure 6C, D**). The percentage of Kv4.2-positive cells was not significantly different between WT and Kv4.3^-/-^ mice (WT 20/423=4.7% *vs* Kv4.3^-/-^ 19/382=5%, p=1, Fisher exact test, **Figure 6D**). Although several proteins, including ion channels (Wolfart et al., 2001; Neuhoff et al., 2002), have been demonstrated to not be homogeneously expressed across the SNc, the percentage of Kv4.2-positive DA neurons was very similar in the medial and lateral SNc of both WT (SNc medial 11/227=4.8% *vs* SNc lateral 9/196=4.6%, p=1, Fisher exact test) and Kv4.3^-/-^ mice (SNc medial 13/218=6% *vs* SNc lateral 6/164=3.7%, p=0.35, Fisher exact test). Thus, these experiments demonstrate that a minority of SNc DA neurons (~5%) displays a strong membrane expression of Kv4.2 and that this pattern of expression is not modified by the loss of Kv4.3. Interestingly, this percentage is not statistically different from the percentage of Kv4.3^-/-^ SNc DA neurons presenting an atypically large and slow I_A_ reported in **Figure 5C-E** (5/42=11.9% vs 19/382=4.6%, p=0.077, Fisher exact test).

**Figure 6.**
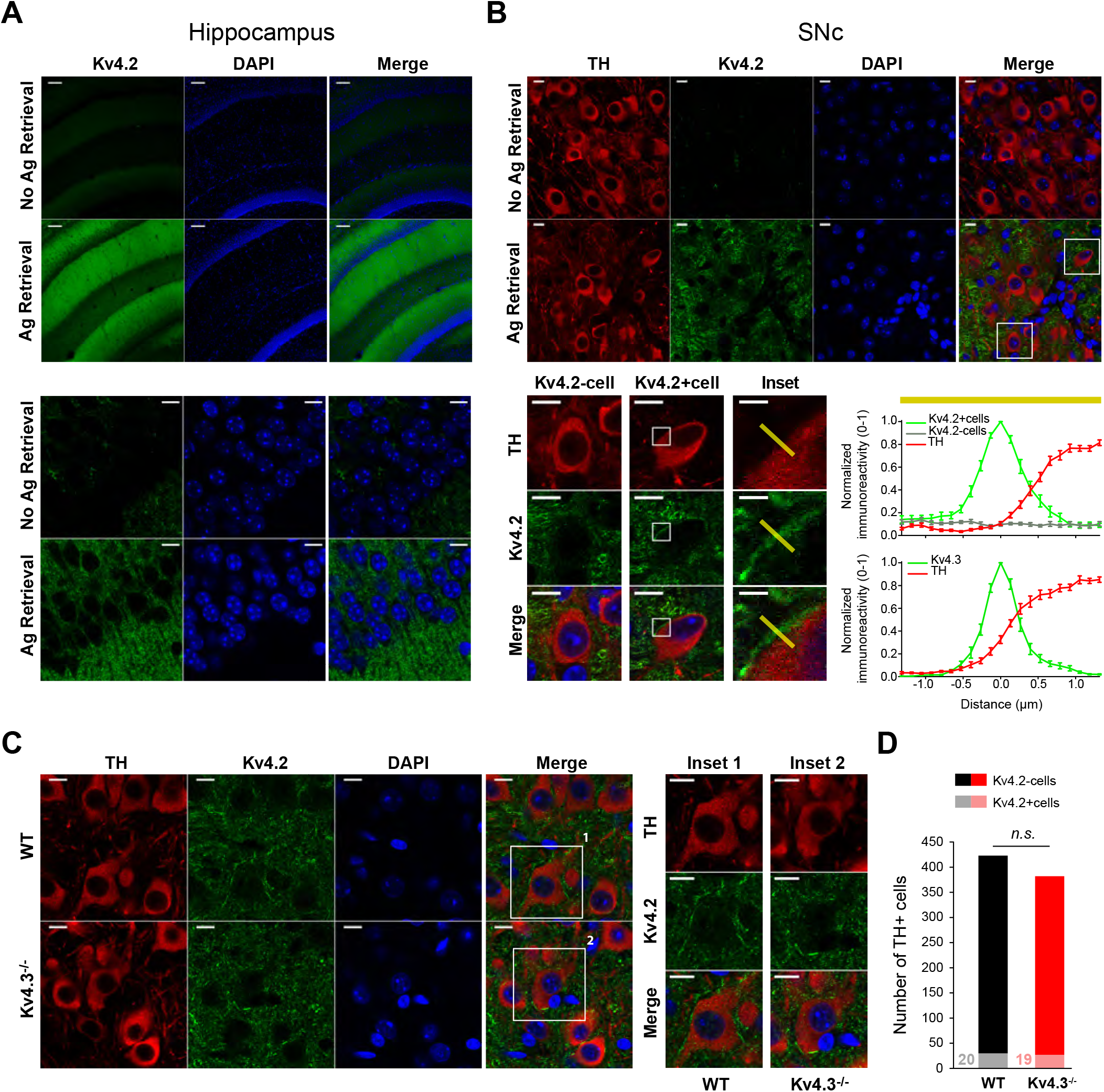
Kv4.2 channels are expressed by a minority of wild-type and Kv4.3^-/-^ SNc DA neurons. **A**, low-magnification (top rows) and high-magnification pictures (bottom rows) showing the effect of the antigen retrieval procedure on Kv4.2 immunostaining (green) in the CA1 region of the hippocampus. DAPI staining (blue) is also shown. **B**, effect of the antigen retrieval procedure on Kv4.2 detection in the SNc. Top, low-magnification pictures showing (from left to right) TH (red), Kv4.2 (green) and DAPI staining (blue) without (top row) or with antigen retrieval (bottom row). Bottom, left, high-magnification pictures of the insets depicted in the merged low-magnification picture and corresponding to a Kv4.2-negative (left) and a Kv4.2-positive SNc DA neuron (middle), for which inset pictures (right) illustrate the region selected to characterize the immunofluorescence profile of Kv4.2 and TH stainings (3μm yellow bar). Bottom, right, immunofluorescence profiles of Kv4.2 (green, top graph), Kv4.3 (green, bottom graph) and TH staining (red, both graphs), showing that both Kv4.2 and Kv4.3 profiles are very similar and strongly suggestive of specific plasma membrane expression. The average profiles (n=5) shown here were defined over 3μm selected regions of (yellow bar above the graph), such as the one shown on the inset pictures on the left. **C**, left, high-magnification pictures showing (from left to right) TH (red), Kv4.2 (green) and DAPI staining (blue) after antigen retrieval in WT (top row) and Kv4.3^-/-^ mice (bottom row). Right, expanded view of the Kv4.2-positive cells (1, 2) highlighted in the merged pictures on the left. **D**, bar plot showing the counts of Kv4.2-negative (dark colors) and -positive (light colors) cells observed in the SNc of WT (black bar) and Kv4.3^-/-^ mice (red bar). n.s. non-significant. Scale bars: **A**, top row 75μm, bottom row 10μm; **B**, top row 10μm, bottom row 10μm, inset 2μm; **C**, 10μm.

### Lack of compensation in the face of Kv4.3 loss

Our results so far suggest that the loss of Kv4.3 is not compensated by changes in expression or function of closely related ion channels. Kv4.2 expression appears to be unchanged in Kv4.3^-/-^ mice (**Figure 6**) and the A-type current is almost completely abolished (10-fold reduction in charge density) in most (~90%) Kv4.3^-/-^ SNc DA neurons (**Figure 5**). In addition, the overall electrophysiological modifications reported (**Figure 3**, **4**) are reminiscent of the reported effects of Kv4-specific toxins on SNc DA neurons (Liss et al., 2001; Amendola et al., 2012; Tarfa et al., 2017). In particular, acute blockade of Kv4 channels is known to induce a strong increase and decrease in pacemaking frequency and post-inhibitory rebound delay, respectively (Liss et al., 2001; Amendola et al., 2012; Tarfa et al., 2017). However, genetic deletion of Kv4.2 channels in cortical pyramidal neurons has been demonstrated to be associated with compensatory modifications in a delayed rectifier-like (IKDR-like) potassium current (Nerbonne et al., 2008). In order to reveal putative homeostatic compensations of Kv4.3 deletion in SNc DA neurons, we first performed a series of current-clamp recordings on a subset of neurons (n=18 for WT, n=32 for Kv4.3^-/-^) to compare the effect of acutely blocking Kv4 channels using the scorpion toxin AmmTX3 (Vacher et al., 2002) to the changes observed in the Kv4.3^-/-^ mouse. Consistent with previous reports (Amendola et al., 2012; Tarfa et al., 2017), AmmTX3 strongly increased pacemaking frequency and dramatically reduced post-inhibitory rebound delay (**Figure 7**). Most interestingly though, the magnitude of the effects of the toxin was very similar to that observed in the Kv4.3^-/-^ neurons: firing frequency was increased by ~68% in both conditions (**Figure 7B**), while rebound delay was decreased by 87% after AmmTX3 and by 82% in Kv4.3^-/-^ mice (**Figure 7D**). Consistent with the data presented earlier, these results strongly suggest that the Kv4-mediated A-type current is virtually completely abolished in Kv4.3^-/-^ SNc DA neurons and that its loss is not compensated by changes in other Kv channels (and associated currents).

**Figure 7.**
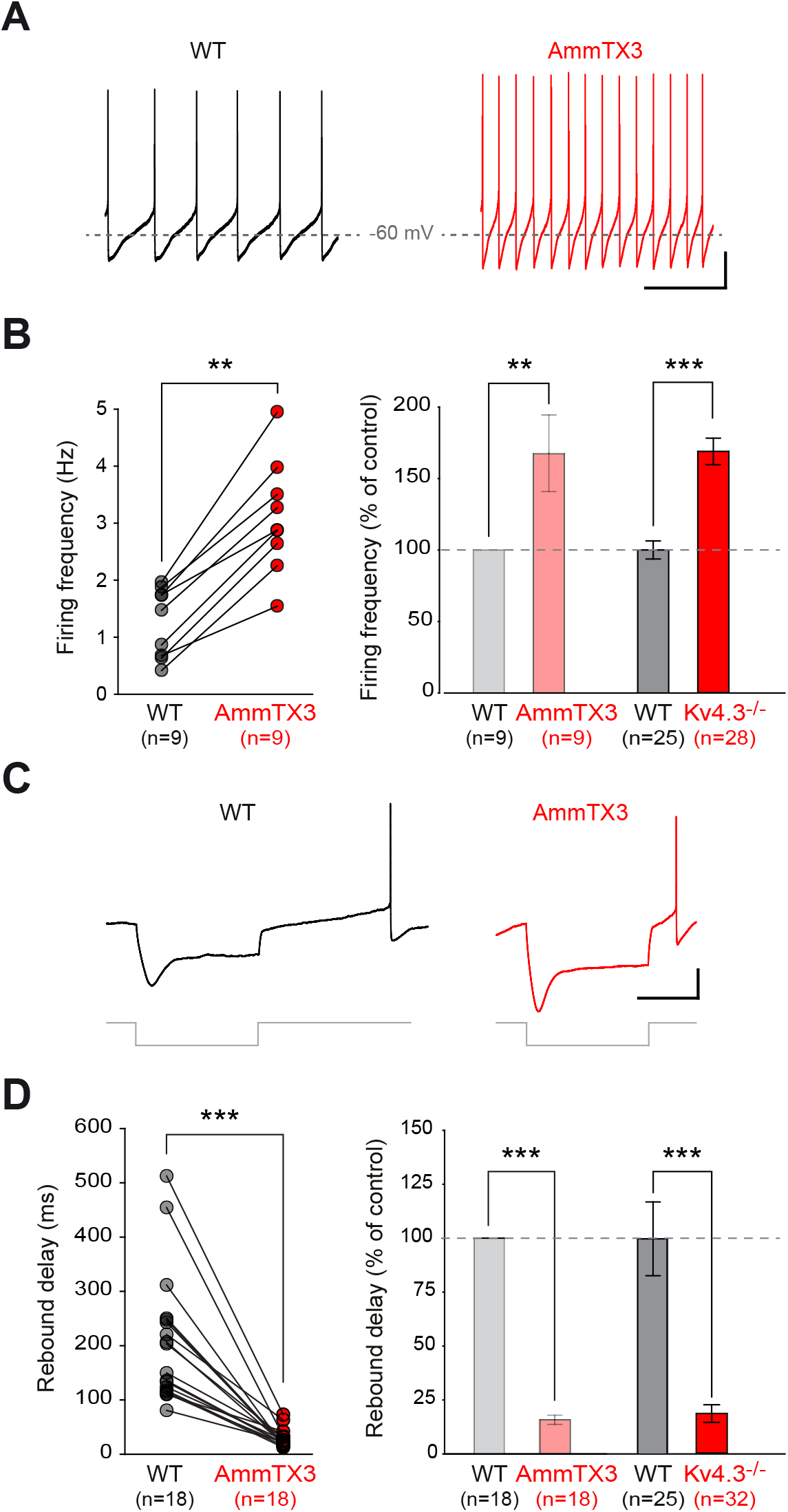
Comparing the alterations in electrophysiological phenotype after acute blockade of Kv4 channels with the Kv4.3^-/-^ mouse model. **A**, current-clamp recordings showing the spontaneous pattern of activity of a WT SNc DA neuron in control condition (black trace, left) and after AmmTX3 application (red trace, right). **B**, left, line and scatter plot showing the change in spontaneous firing frequency induced by AmmTX3 application in individual WT SNc DA neurons. Right, bar plot comparing the average change in spontaneous firing frequency after AmmTX3 application (left, light colors) or Kv4.3 channel deletion (right, dark colors). **C**, current-clamp recordings showing the voltage response of a WT SNc DA neuron to a hyperpolarizing current step (bottom gray traces) in control condition (left, black trace) and after AmmTX3 application (right, red trace). **D**, left, line and scatter plot showing the change in rebound delay induced by AmmTX3 application in individual WT SNc DA neurons. Right, bar plot showing the average change in rebound delay after AmmTX3 application (left, light colors) or Kv4.3 channel deletion (right, dark colors). ** p<0.01, *** p<0.001. Scale bars: **A**, horizontal 1s, vertical 20mV, horizontal gray dotted lines indicate −60mV; **C**, horizontal 500ms, vertical 20mV.

I_H_ and I_A_ have been shown to have an opposite influence on post-inhibitory firing in SNc DA neurons (Amendola et al., 2012; Tarfa et al., 2017), suggesting that a decrease in I_A_ could be compensated by a parallel decrease in I_H_ (Amendola et al., 2012). Although the results presented so far suggest that Kv4.3 loss is not compensated for in SNc DA neurons (**Figures 5**, **6**, **7**), we used voltage-clamp recordings to directly assess whether I_KDR_ (as the results from (Nerbonne et al., 2008) may suggest) and I_H_ could be modified in Kv4.3^-/-^ SNc DA neurons (**Figure 8**). Unlike what has been described in cortical neurons following Kv4.2 deletion, I_KDR_ was not modified in Kv4.3^-/-^ SNc DA neurons (**Figure 8A**). I_H_ was found to be slightly larger in Kv4.3^-/-^ SNc DA neurons (**Figure 8B, Table 2**), but its voltage sensitivity was unchanged. Altogether, the voltage-clamp recordings of I_A_ (**Figure 5**), I_KDR_ and I_H_ (**Figure 8**) and the AR Kv4.2 immunostaining (**Figure 6**) suggest that Kv4.3 loss is not compensated by changes in expression and/or function of functionally-overlapping channels. These data provide a biophysical explanation for the observation made earlier that the acute blockade of Kv4 channels produces an electrophysiological phenotype qualitatively and quantitatively virtually identical to the Kv4.3 genetic deletion (**Figure 7**).

**Figure 8.**
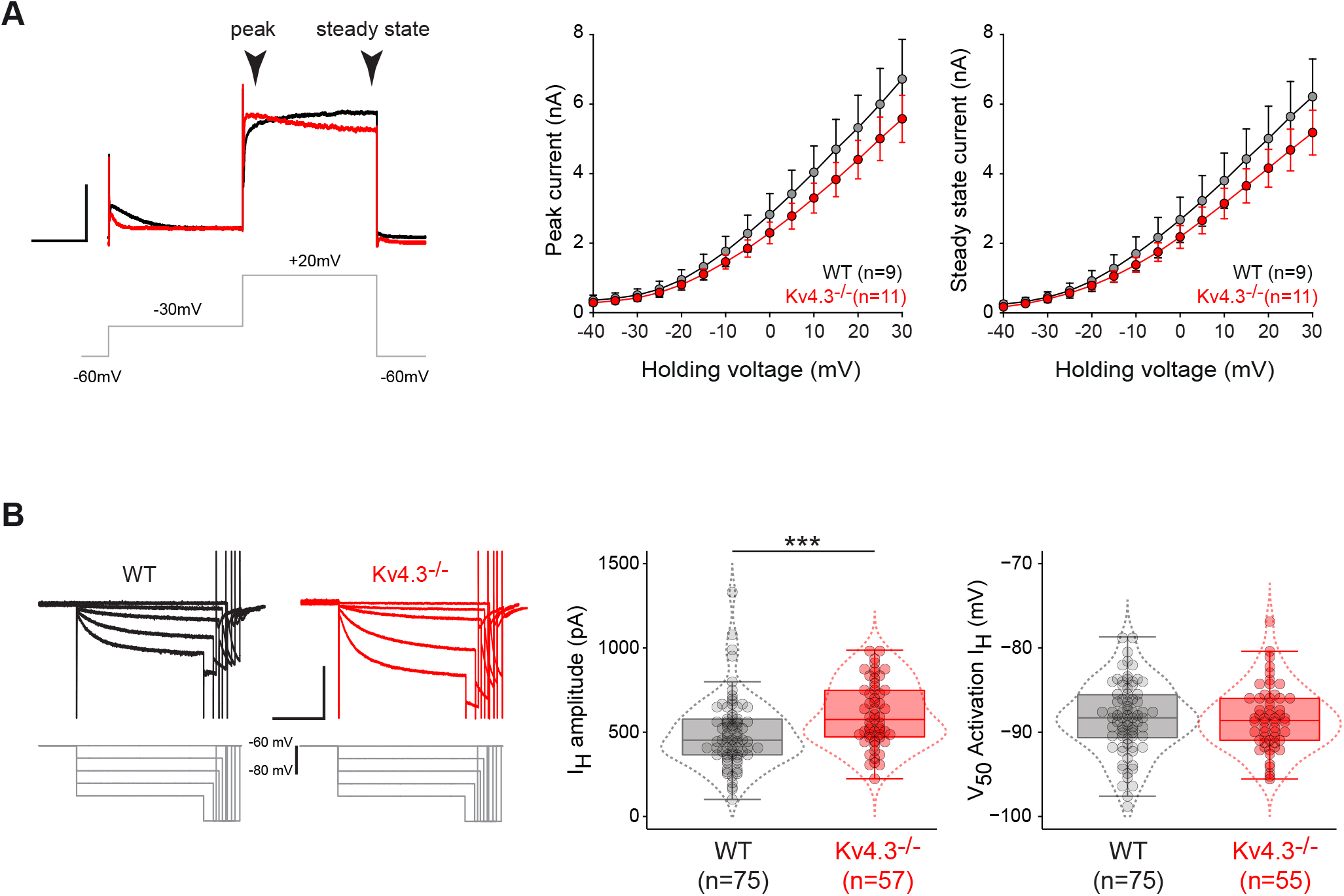
Absence of compensatory changes in delayed rectifier and I_H_ currents in the Kv4.3^-/-^ SNc DA neurons. **A**, properties of the delayed rectifier potassium current (I_KDR_) in SNc DA neurons in WT and Kv4.3^-/-^ mice. Left, voltage-clamp recordings of I_KDR_ obtained in a WT (black trace) and a Kv4.3^-/-^ mouse (red trace) in response to a voltage step to +20mV (gray trace). The peak and steady-state components of IKDR are indicated by arrowheads. Right, line and scatter plots representing the average currentvoltage relationships of the peak (left) and steady-state IKDR (right) obtained from WT (gray dots) and Kv4.3^-/-^ SNc DA neurons (red dots). **B**, properties of I_H_ in SNc DA neurons in WT and Kv4.3^-/-^ mice. Left, voltage-clamp recordings of I_H_ obtained in a WT (black traces) and a Kv4.3^-/-^ mouse (red traces) in response to increasingly hyperpolarized voltage steps. Right, box and whisker plot showing the distribution of values for I_H_ amplitude (left) and voltage sensitivity (right) in WT and Kv4.3^-/-^ SNc DA neurons. *** p<0.001. Dotted lines in the box and whisker plots indicate the distribution of data (violin plots). Scale bars: **A**, horizontal 200ms, vertical 1nA; **B**, horizontal 2s, vertical 500pA.

### Bridging the gap between biophysical changes in I_A_ and I_H_ and variation in electrophysiological phenotype

The computational modeling results presented at the beginning of this study suggested that I_A_ maximal conductance and voltage-dependence (coupled to I_H_ voltage-dependence) were the two biophysical parameters with the greatest influence on SNc DA neuron firing (**Figure 1B, D**). However, an important difference between the model and real SNc DA neurons is that many other conductances may also vary in their expression level and biophysical properties from neuron to neuron, compensating or enhancing the effect of variations in I_A_ or I_H_ properties (Schulz et al., 2006; Goaillard et al., 2009; Amendola et al., 2012). We therefore decided to investigate whether the variations in I_A_ biophysical properties in WT and Kv4.3^-/-^ SNc DA neurons were predictive of variations in electrophysiological phenotype (**Figure 9**). We first looked at potential correlations between firing parameters. As already presented in **Figure 3**, the most significant alterations in firing observed in Kv4.3^-/-^ SNc DA neurons are a strong increase in spontaneous firing frequency (strong decrease in the extracellularly-measured ISI) and a strong decrease in post-inhibitory rebound delay. We found that extracellular ISI and rebound delay (log transformed) were strongly positively correlated with each other in both WT and Kv4.3^-/-^ neurons (**Figure 9B**), although the slope of this relationship may be slightly different between the two genotypes. This correlation is consistent with the reported role of I_A_ in controlling both spontaneous activity and post-inhibitory rebound (Liss et al., 2001; Putzier et al., 2008; Amendola et al., 2012; Tarfa et al., 2017). We therefore tried to determine whether specific I_A_ biophysical properties were better predictors of these variations in ISI or rebound delay (**Figure 9C**). As I_H_ has a complementary influence on rebound and has been described to be co-regulated with I_A_ (Amendola et al., 2012; Tarfa et al., 2017), we also analyzed the relationship between I_H_ properties and ISI or rebound delay. Out of the 5 biophysical parameters analyzed (I_A_ tau, I_H_ amplitude, I_A_ amplitude, I_A_ inactivation V_50_, I_H_ activation V_50_), only two parameters were significantly correlated with ISI or rebound delay: I_A_ tau and I_H_ amplitude (log transformed) were positively and negatively correlated, respectively, with both ISI and rebound delay in both WT and Kv4.3^-/-^ neurons (**Figure 9C**). Surprisingly, neither I_A_ amplitude (measured at −40mV) nor its voltage-dependence (inactivation V_50_) were predictive of variations in ISI or rebound delay (**Figure 9C**). I_H_ activation V_50_ was also unable to predict variations in these firing parameters (data not shown). In addition, consistent with the observation made in rat neurons (Amendola et al., 2012), I_H_ activation and I_A_ inactivation V_50_s were found to be positively correlated (**Figure 9D**). I_A_ tau and I_H_ amplitude were also found to be negatively correlated (**Figure 9D**). Based on these observations, we then sought to determine whether combining several of these 5 biophysical parameters could improve the prediction of ISI or rebound delay using multiple linear regression (**Figure 9E**). Similar to what was performed with the modeling data in **Figure 1D**, we first standardized these parameters (subtracting the mean and dividing by the SD), and then looked for the best subset of variables predictive of ISI or rebound delay. While ISI was best predicted by a multiple linear regression involving only I_A_ tau and I_H_ amplitude, rebound delay was better predicted when I_A_ tau, I_H_ amplitude and I_A_ inactivation V_50_ were included in the linear regression (**Figure 9E**). Consistent with the almost exclusive control of rebound delay by I_A_ and I_H_ (Amendola et al., 2012; Tarfa et al., 2017), rebound delay prediction was much more accurate than ISI prediction (r^2^=0.771 compared to r^2^=0.421). Based on the scaling factors given by the multiple linear regression, it is important to note that both ISI and rebound delay in real neurons seem to be most sensitive to variations in I_A_ tau. While I_H_ amplitude has a quantitatively similar influence on ISI (scaling factor 0.05 compared to 0.063 for I_A_ tau), I_A_ V_50_ and I_H_ amplitude together account for ~1/3 of rebound delay variation (scaling factors 0.06 and 0.048, respectively), ~2/3 being explained by I_A_ tau (scaling factor 0.21). While these results are qualitatively compatible with the predictions made by the model, they provide unique insights into which biophysical properties of I_A_ and I_H_ quantitatively determine the specific influence of these currents on spontaneous activity and rebound delay. Both firing features seem to be strongly controlled by the time constant of I_A_ inactivation, the modification of this biophysical variable being crucial to understand the change in phenotype observed in Kv4.3^-/-^ SNc DA neurons.

**Figure 9.**
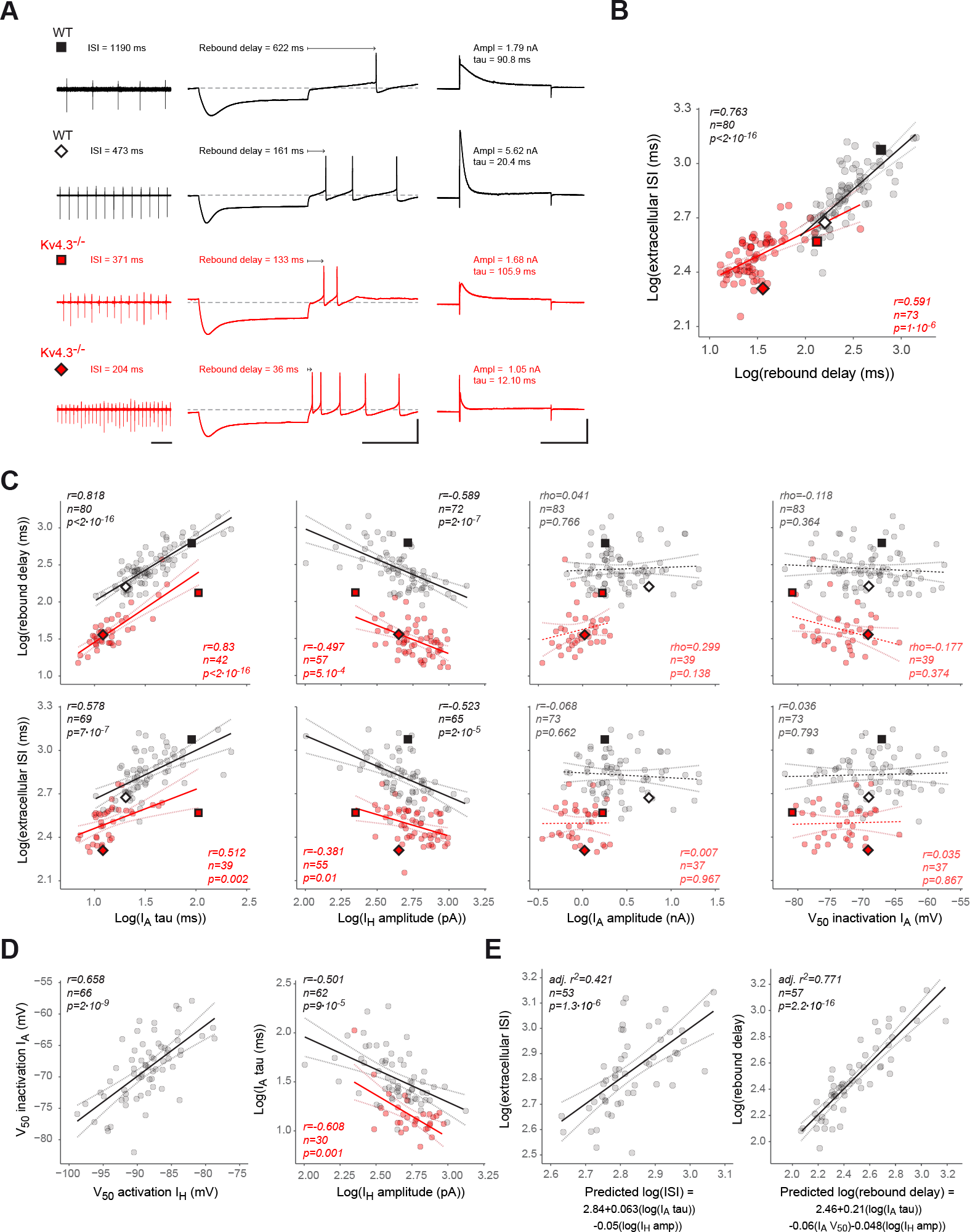
Linking biophysical changes in I_A_ to changes in electrophysiological phenotype in wild-type and Kv4.3^-/-^ SNc DA neurons. **A**, recordings representative of the variation in spontaneous activity, post-inhibitory rebound and I_A_ in WT and Kv4.3’ ‘ SNc DA neurons. Left, cell-attached voltage-clamp recordings of spontaneous pacemaking activity in 2 WT (black traces) and 2 Kv4.3^-/-^ neurons (red traces). The value of the average ISI is indicated above the trace. Middle, current-clamp recordings of the post-inhibitory rebound obtained in the same neurons. The value of rebound delay is indicated above the trace. Right, I_A_ voltage-clamp recordings obtained at −40mV in the same neurons. The values of I_A_ amplitude and tau are indicated above the trace. **B**, scatter plot showing the significant positive correlation between extracellularly recorded ISI and rebound delay observed in WT (gray dots) and Kv4.3^-/-^ SNc DA neurons (red dots). The plain black and red lines correspond to the linear regression of the data (r, n and p values are shown on the graph), while the gray and pink dotted lines indicate the regression confidence intervals. The diamond and square symbols correspond to the recordings presented in panel **A**. **C**, scatter plots showing the relationships between biophysical variables and neuronal output. Correlations between I_A_ tau, I_H_ amplitude, I_A_ amplitude or I_A_ V_50_ (from left to right) and rebound delay or extracellular ISI (from top to bottom) were tested in both WT (gray dots) and Kv4.3^-/-^ neurons (red dots). Please note that only I_A_ tau and I_H_ amplitude were significantly correlated with both ISI and rebound delay in WT and Kv4.3^-/-^ neurons. The plain black and red lines correspond to the linear regression of the data (r/rho, n and p values are shown on the graph), while the gray and pink dotted lines indicate the regression confidence intervals. Dashed black and red lines indicate non-significant correlations. The diamond and square symbols correspond to the recordings presented in panel **A**. **D**, scatter plots showing the significant correlations between I_A_ and I_H_ properties. Left, scatter plot showing the positive correlation between I_A_ inactivation V_50_ and I_H_ activation V_50_ in WT neurons. Right, scatter plot showing the negative correlation between I_A_ tau and I_H_ amplitude observed in both WT (gray dots) and Kv4.3^-/-^ neurons (red dots). The plain black and red lines correspond to the linear regression of the data (r/rho, n and p values are shown on the graph), while the gray and pink dotted lines indicate the regression confidence intervals. **E**, left, scatter plot showing the multiple linear regression of extracellular ISI *vs* I_A_ tau and I_H_ amplitude (predicted ISI) in WT neurons. Right, scatter plot showing the multiple linear regression of rebound delay *vs* I_A_ tau, I_A_ inactivation V_50_ and I_H_ amplitude (predicted rebound delay) in WT neurons. The corresponding equations are indicated below the X axis of each graph. The plain black lines correspond to the linear regression of the data (r, n and p values are shown on the graph), while the gray dotted lines indicate the regression confidence intervals. Scale bars: **A**, left horizontal 1s; middle horizontal 500ms, vertical 50mV; right horizontal 250ms, vertical 2nA.

## DISCUSSION

The current study provides important elements regarding the identity of the ion channels underlying I_A_ in SNc DA neurons and the relative influence of specific biophysical parameters of I_A_ (voltage-dependence, gating kinetics, maximal conductance) on neuronal output. In particular, in contrast with previous studies (Serodio and Rudy, 1998; Liss et al., 2001; Dufour et al., 2014a), we show that the Kv4.2 subunit is expressed in SNc DA neurons, although its functional influence appears negligible in most neurons, due to a very low level of expression. Using Kv4.3^-/-^ mice, we demonstrate that neither the Kv4.2-mediated I_A_, nor the I_KDR_ (delayed rectifier-like) nor I_H_ display biophysical changes compensating for the loss of the Kv4.3-mediated I_A_. Consistent with these findings, the change in electrophysiological phenotype observed in the Kv4.3^-/-^ SNc DA neurons is virtually identical to the one observed after acute blockade of Kv4 channels in WT neurons. In addition, while previous studies (Liss et al., 2001; Putzier et al., 2008) and the computational modeling performed here suggested a strong role of I_A_ conductance in controlling firing frequency, we demonstrate that I_A_ gating kinetics appear as the major determinant of both pacemaking frequency and post-inhibitory rebound delay. Our results also highlight functional complementarity and correlation of biophysical properties of I_A_ and I_H_ in these neurons.

### Kv4.2 is expressed in SNc DA neurons

One of the important results of the present work is the demonstration that Kv4.2 is expressed in mouse SNc DA neurons. So far, it was thought that only Kv4.3 was expressed and entirely responsible for the large A-type current observed in these neurons (Liss and Roeper, 2008; Gantz et al., 2018). Indeed, while single-cell qPCR measurements revealed the presence of Kv4.2 mRNA (Ding et al., 2011; Tapia et al., 2018), classical immunohistochemistry failed to show the presence of the protein subunit (Liss et al., 2001; Dufour et al., 2014a). The use of the Kv4.3^-/-^ mice and antigen retrieval immunohistochemistry however demonstrate that Kv4.2 is expressed by SNc DA neurons. First, in the absence of Kv4.1 (Liss et al., 2001; Ding et al., 2011), the presence of an AmmTX3-sensitive residual A-type current in the Kv4.3^-/-^ neurons is only compatible with the expression of Kv4.2. While the Kv4.2-mediated residual I_A_ is very small and fast in most neurons, in 12% of the voltage clamp-recorded neurons it is large enough to confer a “wild type” phenotype (see **Figure 5, 9**). In the absence of Kv4.3, even though this residual I_A_ current is much faster than its WT counterpart in most neurons, it still influences firing, particularly rebound delay, as suggested by the highly significant correlation between I_A_ tau and rebound delay. Second, antigen retrieval immunohistochemistry confirmed that a Kv4.2 staining is observed in a minority (~5%) of SNc DA neurons in both Kv4.3^-/-^ and WT mice. The perisomatic membranous staining profile, very similar to the one observed for Kv4.3, is consistent with the functional expression of Kv4.2 in these neurons. While the percentage of cells displaying a “high” expression of Kv4.2 is too small to allow a combined voltage-clamp/AR immunohistochemistry approach, the similarity in the proportion of AmmTX3-sensitive “large residual” I_A_ (~12%) and Kv4.2-positive neurons (~5%, not statistically different) strongly suggests that Kv4.2 is responsible for a large I_A_ in a minority of SNc DA neurons. The presence of a small AmmTX3-sensitive residual I_A_ in the rest of the Kv4.3^-/-^ SNc DA neurons suggests that Kv4.2 is expressed and functional in all SNc DA neurons, although its influence might be minor in the presence of Kv4.3. The fact that the residual A-type current in Kv4.3^-/-^ neurons inactivates faster than its WT counterpart is also consistent with the reported differences in inactivation kinetics of Kv4.2 and Kv4.3 (Serodio et al., 1994; Serodio et al., 1996). Altogether, these results suggest that, in contrast to the widely accepted view, a small subpopulation of SNc DA neurons (~5-10%) display an A-type current likely mediated by both Kv4.3 and Kv4.2 channels. In the rest of the SNc DA neurons (~90-95%) however, the contribution of Kv4.2 channels to I_A_ is likely to be negligible, consistent with the phenotype observed in the vast majority of Kv4.3^-/-^ neurons. Interestingly, these two subunits are very similar and coimmunoprecipitate from mouse brain lysates (Marionneau et al., 2009), suggesting that they could form heteromeric I_A_ channels. At this point however, it is difficult to determine whether the Kv4.2-positive neurons in WT mice express Kv4.3/Kv4.2 heteromers or distinct Kv4.3 and Kv4.2 homomers.

### Lack of homeostatic compensation in Kv4.3^-/-^ SNc DA neurons

Intriguingly, these results not only suggest that Kv4.2 expression is very low in most SNc DA neurons but also that it is not modified in the Kv4.3^-/-^ SNc DA neurons. As mentioned before, Kv4.3 and Kv4.2 subunits are very similar. This similarity means that in theory Kv4.2 channels should be able to compensate for the loss of Kv4.3. Despite this, the Kv4.2 pattern of expression is not statistically different between WT and Kv4.3^-/-^ SNc DA neurons and only ~5-10% of the Kv4.3^-/-^ neurons display a “wild type” phenotype. This is reminiscent of previous studies performed on the Kv4.2^-/-^ mouse demonstrating that Kv4.3 expression pattern, assessed by western-blot or immunohistochemistry, is not modified following Kv4.2 loss (Menegola and Trimmer, 2006; Nerbonne et al., 2008). Although the lack of compensation of Kv4.3 loss by Kv4.2 might be surprising, it is important to note that other currents (such as IKDR or I_H_) also appear to not be regulated in a compensatory direction in Kv4.3^-/-^ SNc DA neurons. The current-clamp comparison between the acute blockade of Kv4 channels in WT neurons and the Kv4.3^-/-^ neurons also supports this idea of a lack of compensatory modifications in functionally-overlapping currents. This is surprising in the light of the results obtained on neonatal cortical pyramidal cells (Nerbonne et al., 2008), where Kv4.2 genetic deletion is almost “fully” compensated by an increase in sustained potassium currents. To explain this difference, we may hypothesize that, although the alteration in electrophysiological phenotype observed in Kv4.3^-/-^ neurons is striking (see **Figure 3** in particular), the change in calcium dynamics associated with the elevated spontaneous activity may not be sufficient to trigger homeostatic regulatory mechanisms (O’Leary et al., 2014). Alternatively, modifications in the properties of incoming excitatory or inhibitory synaptic inputs (not analyzed in the current study) may compensate for the changes in intrinsic activity reported here, such that the overall *in vivo* activity of the SNc network is maintained in Kv4.3^-/-^ animals. In line with this hypothesis, Kv4.3^-/-^ mice exhibit only very mild changes in locomotor behavior and motor learning, suggesting that compensatory mechanisms are involved.

### I_A_ gating kinetics play a central role in SNc DA neuron output

The use of I_A_ voltage-clamp measurements and current-clamp recordings on a large number of neurons in WT and Kv4.3^-/-^ mice allowed us to determine the impact of cell-to-cell variations in I_A_ biophysical properties on spontaneous activity and rebound delay. Interestingly, while realistic multi-compartment modeling suggested that I_A_ maximal conductance and voltage-dependence were the two factors most strongly influencing these electrophysiological features, our recordings revealed that I_A_ inactivation rate was the dominant factor defining pacemaking frequency and rebound delay (**Figure 9E**). I_H_ amplitude (proportional to I_H_ maximal conductance in our measurements) was also found to play an important role in real neurons, while its influence was minor in the model. These differences may be explained by several factors. First, the database approach used for our simulations implies that all the tested biophysical properties are varied independently (except for the strict correlation applied to I_A_ and I_H_ V_50_s). While the independence between these biophysical parameters allowed us to precisely quantify the sensitivity of spontaneous activity and rebound delay to each parameter, it does not correspond to the observations made in real neurons: I_H_ amplitude and I_A_ inactivation rate are negatively correlated, i.e. not independent from each other. On the other hand, consistent with the results obtained in rat neurons (Amendola et al., 2012), we demonstrated that I_A_ and I_H_ V_50_s are also positively correlated in WT mouse neurons. However this correlation is far from being perfect (r=0.658, r^2^=0.43). As demonstrated previously (Amendola et al., 2012), the positive correlation in V_50_s enhances the synergistic effect of I_A_ and I_H_ on rebound delay (and likely spontaneous activity), such that applying a strict correlation (r^2^=1) may explain why this biophysical parameter appears as one of the most efficient in modulating firing in the model. These differences in independence of the biophysical properties may partly explain why the model and the experimental observations give different answers. Another factor may explain why I_A_ maximal conductance has a strong effect on firing in the model, but not in real neurons. In order to isolate the effect of I_A_ and I_H_ biophysical properties on firing, all other conductances included in the model were held at fixed values. However, every ion current displays significant cell-to-cell variations in its properties (gating, conductance density) in a same neuronal population (Swensen and Bean, 2005; Schulz et al., 2006; Amendola et al., 2012; Moubarak et al., 2019). If happening at random, these variations in other currents would most likely dampen the effect of the variations in I_A_ or I_H_ specific properties on firing. In fact, we demonstrated in a previous study (Tapia et al., 2018) that the level of expression of Kv4.3 (at the mRNA level) in midbrain DA neurons co-varies with the expression levels of multiple somatodendritic ion channels, including Nav1.2, SK3 and GIRK2. If this co-variation is retained at the protein level, it would mean that cell-to-cell variations in I_A_ maximal conductance occur in parallel with variations in density of other ion channels. By scaling the overall background conductance, these co-variations would also exert a dampening effect on the cell-to-cell variations in I_A_ conductance. Whether correlated or not, cell-to-cell variations in other conductances may thus explain why I_A_ amplitude does not predict pacemaking and rebound delay in real neurons, and why I_A_ inactivation rate appears as the main predictor of these electrophysiological features. In contrast with our findings, the results obtained by Liss and colleagues (2001) suggested that Kv4.3 expression level and channel density predicted pacemaking frequency in mouse neurons. Interestingly, it is noteworthy that I_A_ inactivation rate showed restricted cell-to-cell variations in their recordings (2-fold range). On the other hand, I_A_ charge density showed a 10-fold range of variation, suggesting that most of the variation in I_A_ function was due to variations in I_A_ maximal conductance (Liss et al., 2001). In our recordings however, the levels of variability observed for I_A_ (and I_H_) biophysical properties were rather similar, which led us to apply a 10-fold range to each parameter in our model. Thus, we postulate that the differences in our conclusions may be essentially related to differences in the cell-to-cell variability range of I_A_ and I_H_ biophysical parameters recorded in our samples.

### Functional complementarity and co-regulation of I_A_ and I_H_ in SNc DA neurons

While our results confirm the well-established influence of I_A_ on SNc DA neuron firing (Liss et al., 2001; Gentet and Williams, 2007; Putzier et al., 2008; Amendola et al., 2012; Tarfa et al., 2017), they also emphasize the functional complementarity between I_A_ and I_H_ in these neurons, and reinforce the idea that the channels underlying these currents are co-regulated (Amendola et al., 2012). Indeed, we confirm that I_A_ and I_H_ voltage-dependences are positively correlated and show that I_H_ amplitude and I_A_ inactivation rate are negatively correlated. While we do not have a mechanistic explanation for this latter correlation, these results are reminiscent of the observations made by Tarfa and colleagues on nigrostriatal and mesoaccumbal DA neurons (2017). These authors demonstrated that the longer post-inhibitory delay observed in mesoaccumbal compared to nigrostriatal neurons is explained by a slower I_A_ combined with a smaller I_H_, suggesting a negative correlation between I_A_ inactivation rate and I_H_ amplitude across these two neuronal populations. At the functional level, our results demonstrate these two parameters are the main predictors of the cell-to-cell variations in pacemaking frequency and rebound delay, reinforcing the idea that I_A_ and I_H_ function as a complementary pair of currents tightly controlling post-inhibitory rebound delay in SNc DA neurons (r^2^=0.77). The fact that pacemaking rate is not as accurately predicted by I_A_ and I_H_ properties (r^2^=0.42) is consistent with the documented role of many other conductances and morphological parameters in defining this firing feature (Nedergaard and Greenfield, 1992; Wilson and Callaway, 2000; Wolfart et al., 2001; Liss et al., 2005; Puopolo et al., 2007; Putzier et al., 2009; Gantz et al., 2018; Moubarak et al., 2019).

## Acknowledgements

this work was supported by the European Research Council (Consolidator grant 616827 *CanaloHmics* to J-M.G., supporting A.H-H. and M.T.), the Fondation de France (grant 00076344 to J-M.G. and M.A., supporting A.H-H.), and the Agence Nationale pour la Recherche (ANR Logik ANR-17-CE16-0022, supporting J.R.F.). We thank O. Toutendji for technical assistance.

## REFERENCES

Amendola J, Woodhouse A, Martin-Eauclaire MF, Goaillard JM (2012) Ca(2)(+)/cAMP-sensitive covariation of I(A) and I(H) voltage dependences tunes rebound firing in dopaminergic neurons. J Neurosci 32:2166–2181.

Atherton JF, Bevan MD (2005) Ionic Mechanisms Underlying Autonomous Action Potential Generation in the Somata and Dendrites of GABAergic Substantia Nigra Pars Reticulata Neurons In Vitro. Journal of Neuroscience 25:8272–8281.

Bean BP (2007) The action potential in mammalian central neurons. Nat Rev Neurosci 8:451–465.

Benjamini Y, Hochberg Y (1995) Controlling the False Discovery Rate - a Practical and Powerful Approach to Multiple Testing. J R Stat Soc B 57:289–300.

Carrasquillo Y, Burkhalter A, Nerbonne JM (2012) A-type K+ channels encoded by Kv4.2, Kv4.3 and Kv1.4 differentially regulate intrinsic excitability of cortical pyramidal neurons. J Physiol 590:3877–3890.

Cembrowski MS, Bachman JL, Wang L, Sugino K, Shields BC, Spruston N (2016) Spatial GeneExpression Gradients Underlie Prominent Heterogeneity of CA1 Pyramidal Neurons. Neuron 89:351–368.

Chan CS, Guzman JN, Ilijic E, Mercer JN, Rick C, Tkatch T, Meredith GE, Surmeier DJ (2007) ‘Rejuvenation’ protects neurons in mouse models of Parkinson’s disease. Nature 447:1081–1086.

Destexhe A, Babloyantz A, Sejnowski TJ (1993) Ionic Mechanisms for Intrinsic Slow Oscillations in Thalamic Relay Neurons. Biophysical Journal 65:1538–1552.

Ding S, Matta SG, Zhou FM (2011) Kv3-like potassium channels are required for sustained high-frequency firing in basal ganglia output neurons. J Neurophysiol 105:554–570.

Dufour MA, Woodhouse A, Goaillard JM (2014a) Somatodendritic ion channel expression in substantia nigra pars compacta dopaminergic neurons across postnatal development. J Neurosci Res 92:981–999.

Dufour MA, Woodhouse A, Amendola J, Goaillard JM (2014b) Non-Linear Developmental Trajectory of Electrical Phenotype in Rat Substantia Nigra Pars Compacta Dopaminergic Neurons. eLife 3:e04059.

Fuzik J, Zeisel A, Mate Z, Calvigioni D, Yanagawa Y, Szabo G, Linnarsson S, Harkany T (2016) Integration of electrophysiological recordings with single-cell RNA-seq data identifies neuronal subtypes. Nat Biotechnol 34:175–183.

Gantz SC, Ford CP, Morikawa H, Williams JT (2018) The Evolving Understanding of Dopamine Neurons in the Substantia Nigra and Ventral Tegmental Area. Annu Rev Physiol 80:219–241.

Gentet LJ, Williams SR (2007) Dopamine gates action potential backpropagation in midbrain dopaminergic neurons. J Neurosci 27:1892–1901.

Goaillard JM, Taylor AL, Schulz DJ, Marder E (2009) Functional consequences of animal-to-animal variation in circuit parameters. Nat Neurosci 12:1424–1430.

Grace AA, Onn SP (1989) Morphology and electrophysiological properties of immunocytochemically identified rat dopamine neurons recorded in vitro. J Neurosci 9:3463–3481.

Guzman JN, Sanchez-Padilla J, Chan CS, Surmeier DJ (2009) Robust pacemaking in substantia nigra dopaminergic neurons. J Neurosci 29:11011–11019.

Hines ML, Carnevale NT (2001) NEURON: A tool for neuroscientists. Neuroscientist 7:123–135.

Hodgkin AL, Huxley AF (1952) A quantitative description of membrane current and its application to conduction and excitation in nerve. J Physiol 117:500–544.

Hu W, Tian C, Li T, Yang M, Hou H, Shu Y (2009) Distinct contributions of Na(v)1.6 and Na(v)1.2 in action potential initiation and backpropagation. Nat Neurosci 12:996–1002.

Ji H, Tucker KR, Putzier I, Huertas MA, Horn JP, Canavier CC, Levitan ES, Shepard PD (2012) Functional characterization of ether-a-go-go-related gene potassium channels in midbrain dopamine neurons - implications for a role in depolarization block. Eur J Neurosci 36:2906–2916.

Jiao Y, Sun ZQ, Lee T, Fusco FR, Kimble TD, Meade CA, Cuthbertson S, Reiner A (1999) A simple and sensitive antigen retrieval method for free-floating and slide-mounted tissue sections. J Neurosci Meth 93:149–162.

Kole MHP, Stuart GJ (2008) Is action potential threshold lowest in the axon? Nat Neurosci 11:1253–1255.

Liss B, Roeper J (2008) Individual dopamine midbrain neurons: functional diversity and flexibility in health and disease. Brain Res Rev 58:314–321.

Liss B, Franz O, Sewing S, Bruns R, Neuhoff H, Roeper J (2001) Tuning pacemaker frequency of individual dopaminergic neurons by Kv4.3L and KChip3.1 transcription. EMBO J 20:5715–5724.

Liss B, Haeckel O, Wildmann J, Miki T, Seino S, Roeper J (2005) K-ATP channels promote the differential degeneration of dopaminergic midbrain neurons. Nat Neurosci 8:1742–1751.

Lorincz A, Nusser Z (2008) Cell-Type-Dependent Molecular Composition of the Axon Initial Segment. J Neurosci 28:14329–14340.

Marionneau C, LeDuc RD, Rohrs HW, Link AJ, Townsend RR, Nerbonne JM (2009) Proteomic analyses of native brain K(V)4.2 channel complexes. Channels (Austin) 3:284–294.

Menegola M, Trimmer JS (2006) Unanticipated region- and cell-specific downregulation of individual KChIP auxiliary subunit isotypes in Kv4.2 knock-out mouse brain. J Neurosci 26:12137–12142.

Moubarak E, Engel D, Dufour MA, Tapia M, Tell F, Goaillard JM (2019) Robustness to Axon Initial Segment Variation Is Explained by Somatodendritic Excitability in Rat Substantia Nigra Dopaminergic Neurons. J Neurosci 39:5044–5063.

Nedergaard S, Greenfield SA (1992) Subpopulations of Pars Compacta Neurons in the Substantia-Nigra - the Significance of Qualitatively and Quantitatively Distinct Conductances. Neuroscience 48:423–437.

Nerbonne JM, Gerber BR, Norris A, Burkhalter A (2008) Electrical remodelling maintains firing properties in cortical pyramidal neurons lacking KCND2-encoded A-type K+ currents. J Physiol 586:1565–1579.

Neuhoff H, Neu A, Liss B, Roeper J (2002) I(h) channels contribute to the different functional properties of identified dopaminergic subpopulations in the midbrain. J Neurosci 22:1290–1302.

Niwa N, Wang W, Sha Q, Marionneau C, Nerbonne JM (2008) Kv4.3 is not required for the generation of functional Ito,f channels in adult mouse ventricles. J Mol Cell Cardiol 44:95–104.

Northcutt AJ, Kick DR, Otopalik AG, Goetz BM, Harris RM, Santin JM, Hofmann HA, Marder E, Schulz DJ (2019) Molecular profiling of single neurons of known identity in two ganglia from the crab Cancer borealis. P Natl Acad Sci USA 116:26980–26990.

O’Leary T, Williams AH, Franci A, Marder E (2014) Cell types, network homeostasis, and pathological compensation from a biologically plausible ion channel expression model. Neuron 82:809–821.

Philippart F, Destreel G, Merino-Sepulveda P, Henny P, Engel D, Seutin V (2016) Differential Somatic Ca2+ Channel Profile in Midbrain Dopaminergic Neurons. J Neurosci 36:7234–7245.

Puopolo M, Raviola E, Bean BP (2007) Roles of subthreshold calcium current and sodium current in spontaneous firing of mouse midbrain dopamine neurons. J Neurosci 27:645–656.

Putzier I, Kullmann PHM, Horn JP, Levitan ES (2008) Dopamine Neuron Responses Depend Exponentially on Pacemaker Interval. J Neurophysiol 101:926–933.

Putzier I, Kullmann PHM, Horn JP, Levitan ES (2009) Cav1.3 Channel Voltage Dependence, Not Ca2+ Selectivity, Drives Pacemaker Activity and Amplifies Bursts in Nigral Dopamine Neurons. J Neurosci 29:15414–15419.

Schulz DJ, Goaillard JM, Marder E (2006) Variable channel expression in identified single and electrically coupled neurons in different animals. Nat Neurosci 9:356–362.

Serodio P, Rudy B (1998) Differential expression of Kv4 K+ channel subunits mediating subthreshold transient K+ (A-type) currents in rat brain. J Neurophysiol 79:1081–1091.

Serodio P, Kentros C, Rudy B (1994) Identification of Molecular-Components of a-Type Channels Activating at Subthreshold Potentials. J Neurophysiol 72:1516–1529.

Serodio P, Vega-Saenz de Miera E, Rudy B (1996) Cloning of a novel component of A-type K+ channels operating at subthreshold potentials with unique expression in heart and brain. J Neurophysiol 75:2174–2179.

Seutin V, Engel D (2010) Differences in Na+ conductance density and Na+ channel functional properties between dopamine and GABA neurons of the rat substantia nigra. J Neurophysiol 103:3099–3114.

Seutin V, Massotte L, Renette MF, Dresse A (2001) Evidence for a modulatory role of Ih on the firing of a subgroup of midbrain dopamine neurons. Neuroreport 12:255–258.

Swensen AM, Bean BP (2005) Robustness of burst firing in dissociated purkinje neurons with acute or long-term reductions in sodium conductance. J Neurosci 25:3509–3520.

Tapia M, Baudot P, Formisano-Treziny C, Dufour MA, Temporal S, Lasserre M, Marqueze-Pouey B, Gabert J, Kobayashi K, Goaillard JM (2018) Neurotransmitter identity and electrophysiological phenotype are genetically coupled in midbrain dopaminergic neurons. Sci Rep 8:13637.

Tarfa RA, Evans RC, Khaliq ZM (2017) Enhanced Sensitivity to Hyperpolarizing Inhibition in Mesoaccumbal Relative to Nigrostriatal Dopamine Neuron Subpopulations. J Neurosci 37:3311–3330.

Taylor AL, Hickey TJ, Prinz AA, Marder E (2006) Structure and visualization of highdimensional conductance spaces. J Neurophysiol 96:891–905.

Vacher H, Alami M, Crest M, Possani LD, Bougis PE, Martin-Eauclaire MF (2002) Expanding the scorpion toxin alpha-KTX 15 family with AmmTX3 from Androctonus mauretanicus. Eur J Biochem 269:6037–6041.

Vandecasteele M, Deniau JM, Venance L (2011) Spike frequency adaptation is developmentally regulated in substantia nigra pars compacta dopaminergic neurons. Neuroscience 192:1–10.

Wilson CJ, Callaway JC (2000) Coupled oscillator model of the dopaminergic neuron of the substantia nigra. J Neurophysiol 83:3084–3100.

Wolfart J, Neuhoff H, Franz O, Roeper J (2001) Differential expression of the small-conductance, calcium-activated potassium channel SK3 is critical for pacemaker control in dopaminergic midbrain neurons. J Neurosci 21:3443–3456.

